# The evolution of Sox gene repertoires and regulation of segmentation in arachnids

**DOI:** 10.1101/2020.06.04.133389

**Authors:** Luis Baudouin-Gonzalez, Anna Schoenauer, Amber Harper, Grace Blakeley, Michael Seiter, Saad Arif, Lauren Sumner-Rooney, Steven Russell, Prashant P. Sharma, Alistair P. McGregor

## Abstract

The Sox family of transcription factors regulate many processes during metazoan development, including stem cell maintenance and nervous system specification. Characterising the repertoires and roles of these genes can therefore provide important insights into animal evolution and development. We further characterised the Sox repertoires of several arachnid species with and without an ancestral whole genome duplication (WGD), and compared their expression between the spider *Parasteatoda tepidariorum* and the harvestman *Phalangium opilio*. We also found that most Sox families have been retained as ohnologs after WGD and evidence for potential subfunctionalisation and/or neofunctionalization events. Our results also suggest that *Sox21b-1* likely regulated segmentation ancestrally in arachnids, playing a similar role to the closely related SoxB gene, *Dichaete*, in insects. We previously showed that *Sox21b-1* is required for the simultaneous formation of prosomal segments and sequential addition of opisthosomal segments in *P. tepidariorum*. We studied the expression and function of *Sox21b-1* further in this spider and found that while this gene regulates the generation of both prosomal and opisthosomal segments, it plays different roles in the formation of these tagmata reflecting their contrasting modes of segmentation and deployment of gene regulatory networks with different architectures.

## Introduction

The Sox (Sry-Related High-Mobility Group box) genes encode an ancient family of transcription factors that play important roles in the regulation of many aspects of animal development (Kamachi and Kondoh 2013). There are eight known groups (A-H) of Sox genes defined by the sequence of their HMG domains (Bowles, et al. 2000; Heenan, et al. 2016). While group A (represented by Sry) is restricted to eutherian mammals and groups G and H are also thought to be lineage specific, representatives of groups B-F are generally found in all metazoan lineages although there have been lineage specific losses (Bowles, et al. 2000). Our work, and that of others, indicate that groups B-F were all represented in the common ancestor of arthropods with each group represented in *Drosophila melanogaster,* for example (Janssen, et al. 2018; Paese, et al. 2018a; Phochanukul and Russell 2010). However, there has been duplication of some groups in other arthropod lineages (Paese, et al. 2018a) and we have shown that the group B genes *Sox21a* and *Sox21b*, as well as the group C, D, E and F genes are represented by at least two genes in the spider *Parasteatoda tepidariorum*. This is consistent with the whole genome duplication (WGD) that took place in an ancestor of arachnopulmonates (Leite, et al. 2018; Schwager, et al. 2017) (fig. 1) and patterns of gene retention observed after independent WGD in vertebrates (Bowles, et al. 2000; Wegner 1999). Further characterisation of the repertoire and role of Sox genes among arachnids has great potential to better understand the evolution of these genes, and their expression and function in the development of arthropods, as well as gene fate and the outcomes of WGD more generally.

**Figure 1.**
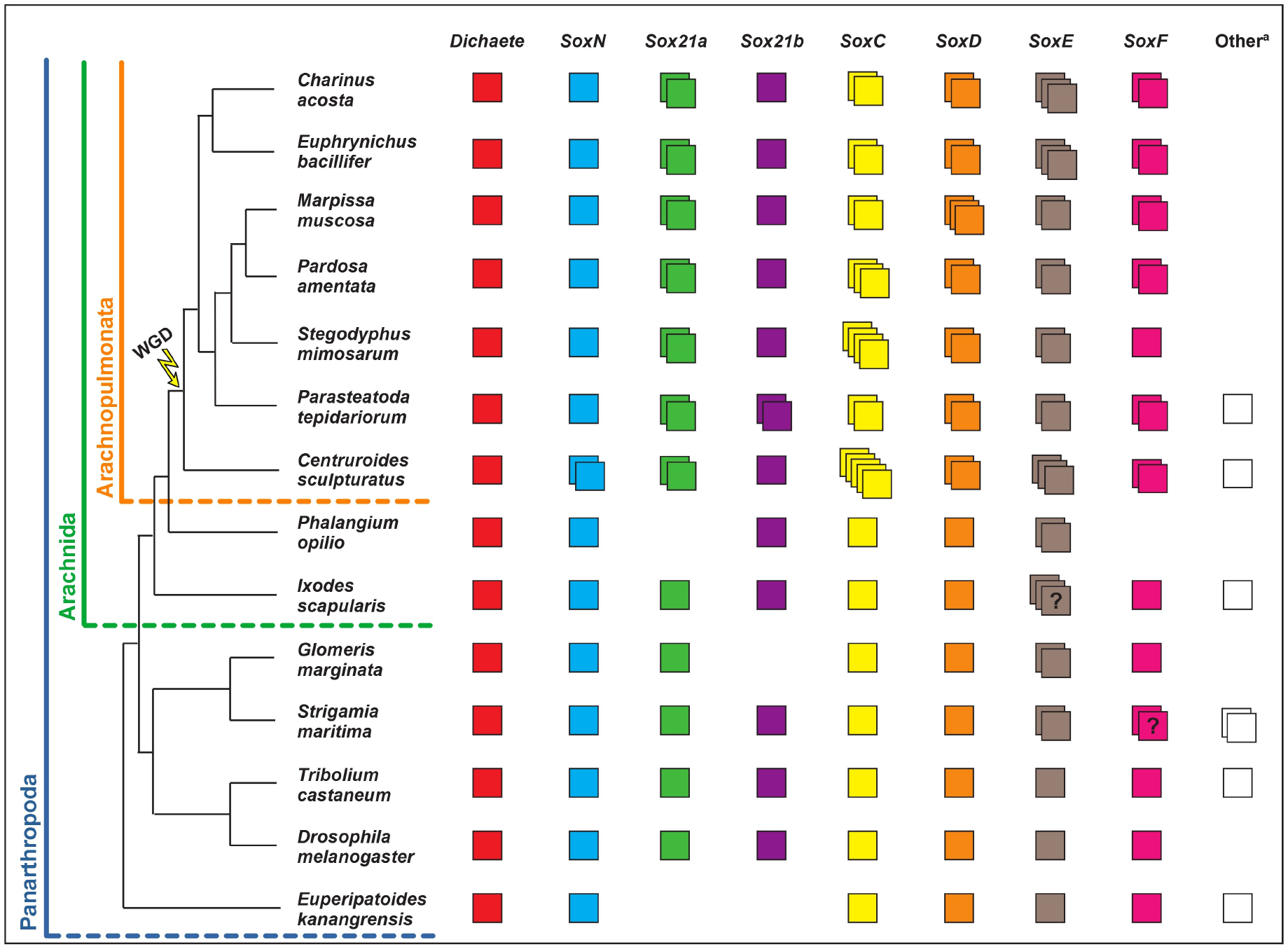
Panarthropod Sox gene repertoires. Sox gene repertoires in the surveyed species (*C. sculpturatus*, *P. opilio*, *I. scapularis*, *S. maritima*, *M. muscosa*, *P. amentata*, *C. acosta* and *E. bacillifer*) and other panarthropods for which the Sox gene repertoires were previously identified (*P. tepidariorum*, *S. mimosarum*, *G. marginata*, *T. castaneum*, *D. melanogaster* and *E. kanangrensis*). Each coloured box represents a different gene.? = unclear number of copies due to incomplete sequences. Other^a^ = unresolved or highly divergent sequences.

We previously found that one of the *P. tepidariorum* SoxB genes, *Sox21b-1,* regulates the formation of segments in this spider (Paese, et al. 2018a; Paese, et al. 2018b). In spiders there are at least two modes of trunk segmentation: the leg bearing segments of the prosoma (cephalothorax) are generated more or less simultaneously like the segments of long germ insects. Opisthosomal (abdominal) segments are added sequentially like most other arthropods, although one at a time instead of two, in contrast to short germ insects (reviewed in Clark, et al. 2019; Hilbrant, et al. 2012; Oda and Akiyama-Oda 2020). It also appears that different gene regulatory networks underlie the simultaneous formation of anterior versus sequential addition of posterior segments (Akiyama-Oda and Oda 2020; McGregor, et al. 2008; Oda, et al. 2007; Paese, et al. 2018b; Pechmann, et al. 2011; Schönauer, et al. 2016; Schwager, et al. 2009). The leg-bearing segments of the prosoma are added through splitting of wider gene expression domains regulated by the gap gene-like activity of *hunchback* (*hb*) and *Distal-less* (*Dll*) (Pechmann, et al. 2011; Schwager, et al. 2009). The opisthosomal segments are sequentially added from a posterior segment addition zone (SAZ), which is in part established by the activity of Wnt and Notch-Delta signalling pathways (Oda, et al. 2007; Schönauer, et al. 2016). The interplay between these pathways directs sequential segment addition from the SAZ by regulating dynamic expression of pair-rule gene orthologs, including *even-skipped* (*eve*), and other segmentation genes such as *caudal* (*cad*) (McGregor, et al. 2008; Oda, et al. 2007; Schönauer, et al. 2016). Knockdown of *Pt-Sox21b-1* expression results in the loss of all leg-bearing prosomal segments and disrupts the formation of the (SAZ), leading to loss of all opisthosomal segments (Paese, et al. 2018b). Therefore *Sox21b-1*, along with *arrow* and *Msx1*, is one of the few genes known to regulate both simultaneous and sequential segmentation in *P. tepidariorum* (Akiyama-Oda and Oda 2020; Paese, et al. 2018b; Setton and Sharma 2018).

Interestingly, *Dichaete,* a closely related SoxB group gene, is found to regulate segmentation in both *D. melanogaster* and *Tribolium castaneum*, long germ and short germ insects respectively (Clark and Peel 2018; Russell, et al. 1996). This finding, together with our work, suggests that SoxB genes regulated segmentation ancestrally in arthropods and continue to do so in simultaneous and sequential segment formation in different lineages.

This conclusion requires a broader understanding of the evolution and expression of Sox genes in other arachnids, as well as a more detailed understanding of the role of these genes, including *Sox21b-1*, in *P. tepidariorum*. To address this, we explored the evolution and roles of Sox genes in arachnid development further by characterising the repertoire of Sox genes in five additional arachnids with and without an ancestral WGD, and compared the expression of Sox genes between the harvestman *Phalangium opilio* and the spider *P. tepidariorum*. Furthermore, we assessed the potential roles of *Dichaete*, *Sox21a-1*, *Sox21b-2* and *SoxD-2* in segmentation in *P. tepidariorum* and examined the expression and function of *Sox21b-1* in more detail. We found similar patterns of retention of duplicated Sox genes after WGD in spiders, scorpions and whip-spiders and evidence for potential sub and/or neofunctionalisation during embryogenesis in the spider *P. tepidariorum*. Our results also suggest that although other Sox genes, including *Dichaete* may not be required for segmentation in *P. tepidariorum*, *Sox21b-1* regulated prosomal and opisthosomal segmentation ancestrally in arachnids as a key component of the gene regulatory networks for their simultaneous versus sequential production, respectively.

## Results

### Arachnid Sox gene repertoires

To further investigate the evolution of the Sox gene repertoires of arachnids compared to other arthropods more broadly, we surveyed new and available genomic resources to identify Sox HMG-domain containing sequences from several additional arachnids: the spiders *Marpissa muscosa* and *Pardosa amentata*, the amblypygids *Charinus acosta* and *Euphrynichus bacillifer*, the scorpion *Centruroides sculpturatus*, the harvestmen *P. opilio* and the tick *Ixodes scapularis*, and a myriapod, the centipede *Strigamia maritima*. Sequences were annotated as the best hits obtained from the SMART BLAST NCBI online tool and by reciprocal BLAST against the proteome of *P. tepidariorum* and compared to previously identified arthropod Sox genes (Janssen, et al. 2018; Paese, et al. 2018a).

Our previous analysis of the *P. tepidariorum* genome identified fifteen Sox genes (Paese et al., 2018b) (fig. 1). We identified fourteen Sox genes in *P. amentata*, fourteen in *M. muscosa*, fourteen in *C. acosta,* fourteen in *E. bacillifer*, nineteen in *C. sculpturatus*, seven in *P. opilio*, eleven in *I. scapularis* and twelve in *S. maritima* (fig. 1 and supplementary file 1, Supplementary Material online). However, some of the sequences retrieved had incomplete HMG domains (10/105), probably representing fragments of the full sequence (supplementary file 2, Supplementary Material online). Therefore, the Sox repertoires described here may be incomplete due to the quality of genomic or transcriptomic assemblies or some genes were potentially not captured in the transcriptomes sequenced, as we found previously for Hox genes (Leite, et al. 2018), because Sox gene loss is very rare, especially in the arthropods (Phochanukul and Russell 2010).

The sequences we obtained were assigned to particular Sox groups based on the best match from reciprocal BLAST and verified using maximum likelihood trees (supplementary figs 1 and 2, Supplementary Material online). The trees supported the classification of most sequences obtained, forming monophyletic clades corresponding to each Sox group (supplementary figs 1 and 2, Supplementary Material online). When constructing trees with all HMG sequences the bootstrap support values were low, as previously found with this domain (Paese, et al. 2018a). However, removing *Pt-SoxB-like*, *Cscu-SoxB-like*, *Isca-SoxB-like*, *Smar-SoxB-like*, *Smar-SoxE-like* and *Tcas-SoxB5*, as well as *Gmar-Sox21a* (which has an incomplete HMG domain sequence) gave better support values for Sox Group B (81) and Sox Group E (80), with Sox Groups D and F still being very well supported (100 and 98 respectively) (supplementary fig. 2, Supplementary Material online). The only exception was Sox Group C genes (34), possibly due to considerable sequence divergence within this group. The monophyly of Sox Groups E and F was also well supported (80), and within Sox Group B, there was strong support for the monophyly of *SoxN* sequences (79), as well as reasonable support for the monophyly of arachnid *Dichaete* sequences (61) and arachnid *Sox21b* sequences (63) (supplementary fig. 2, Supplementary Material online). Together, these analyses are concordant with the phylogenetic relationships for the Sox family previously established using vertebrate and invertebrate Sox genes (Bowles, et al. 2000; McKimmie, et al. 2005; Wilson and Dearden 2008).

At least one representative of each Sox group was found for all species surveyed, with the exception of *Sox21a* and *SoxF* in *P. opilio* (fig. 1). Consistent with previous surveys (Janssen, et al. 2018; Paese, et al. 2018a), a single copy of *Dichaete* was found in all species analysed (fig. 1). A single copy of *SoxN* was also found in most species, with the exception of *C. sculpturatus* where two sequences with identical HMG domains were found, although they differ outside this domain (fig. 1). A single copy of *Sox21a* is present in non-WGD arthropods, whereas two copies were found in all seven arachnopulmonate species analysed (fig. 1). Additional *SoxB-like* sequences were also found in *C. sculpturatus*, *I. scapularis* and *S. maritima*, although, much like other previously identified *SoxB-like* genes (i.e. *Tcas-SoxB5* and *Ekan-SoxB3*) (Janssen, et al. 2018; Paese, et al. 2018a), these sequences diverge significantly from other Group B genes and therefore cannot be reliably classified (fig. 1 and supplementary fig. 2, Supplementary Material online).

All non-WGD arthropods (insects, myriapod, harvestman and tick) have single copies of *SoxC* and *SoxD*, but in arachnopulmonates, at least two copies for each of these groups were found (fig. 1). At least two *SoxE* and *SoxF* genes were also found in all arachnopulmonates except for *S. mimosarum,* where *SoxF* is only represented by one gene, and in *P. opilio SoxF* is potentially missing (fig. 1). *SoxE* also appears to be duplicated in *P. opilio, G. marginata*, *S. maritima* and *I. scapularis*, although *Smar-SoxE2* has an incomplete domain, and two of the three copies in *I. scapularis* contain non-overlapping fragments of the HMG domain (indicating they might be the same gene) (fig. 1 and supplementary fig. 2, Supplementary Material online). An additional *SoxE-like* sequence was found in *S. maritima*, although this sequence is highly divergent from other *SoxE* genes (fig. 1 and supplementary fig. 2, Supplementary Material online). We note that *SoxE* duplication may be relatively common in invertebrate lineages compared to other Sox families, with duplications previously identified in Hymenoptera (Wilson and Dearden 2008). Two *SoxF* genes were also found in *S. maritima* but they have incomplete HMG domains, and so may represent fragments of a single gene (fig. 1 and supplementary fig. 2, Supplementary Material online).

Finally, our survey revealed an interesting evolutionary pattern for *Sox21b*. It appears that among the species we have surveyed this Sox gene is only duplicated in *P. tepidariorum* (fig. 1). This suggests that after the arachnopulmonate WGD both ohnologs of *Sox21b* were only retained in a restricted lineage including *P. tepidariorum* or that this Sox gene was tandemly duplicated later in this spider, which raises many interesting questions regarding the evolution of its role during segmentation in arachnids.

### Expression of Sox genes during embryogenesis in P. tepidariorum and P. opilio

To help better understand the evolution and roles of Sox genes in arachnids, we compared the expression patterns of the *P. tepidariorum* Sox genes with their orthologs in *P. opilio*. Although we previously characterised Sox gene expression patterns in *P. tepidariorum*, we only detected expression for six out of 15 Sox genes present in this species (*Pt-SoxN*, *Pt-Sox21b-1*, *Pt-SoxC1*, *Pt-SoxD1*, *Pt-SoxE1* and *Pt-SoxF2*) (Paese, et al. 2018a). However, it remained likely that the other genes were expressed at least at low levels during embryogenesis because their transcripts could be detected in RNA-Seq data (Iwasaki-Yokozawa, et al. 2018). We therefore performed additional analyses using longer in situ hybridisation probes on a wider range of embryonic stages.

The expression patterns observed for the *P. tepidariorum* Sox genes were generally the same as those we reported previously (Paese, et al. 2018a) (supplementary figs 3-9, Supplementary Material online). However, we found that *Pt-D* is expressed in a diffuse salt and pepper pattern in the developing cephalic lobe and prosoma at stages 6 and 7, as well as potentially in the SAZ (although this may be background signal) (supplementary fig. 3, Supplementary Material online). Subsequently we detected *Pt-D* in segmental stripes in L1 to L3 at stages 8.1 and 8.2, as well as along the ventral midline and in the prosomal appendages at stage 10.1 (supplementary fig. 3, Supplementary Material online). In addition, we were now able to detect expression of *Pt-Sox21a-1* and *Pt-Sox21a-2* in non-overlapping patterns mainly in the neuroectoderm, which is suggestive of sub- and/or neofunctionalisation (supplementary fig. 5, Supplementary Material online). Furthermore, *Pt-Sox21a-1* appears to be expressed in segmental stripes in L1 and L2 at stage 8.1 and more ventrally restricted stripes in the opisthosoma at stage 9.2, which suggests it may play a role in segmentation (supplementary fig. 5, Supplementary Material online). We also observed *Pt-SoxE-2* expression in a similar pattern to *Pt-SoxE-1* in the prosomal and opisthosomal limb buds, albeit with some temporal and spatial differences, as well as unique expression of *Pt-SoxE-2* in opisthosomal cells that are possibly the germline progenitors (Schwager, et al. 2015) (supplementary fig. 8, Supplementary Material online). We also found that while *Pt-SoxF-1* and *Pt-SoxF-2* are both expressed in the developing prosomal and opisthosomal appendages, each has also a specific domain of expression: *Pt-SoxF-1* is expressed in the secondary eye primordia (supplementary fig. 9, Supplementary Material online), and *Pt-SoxF-2* is expressed along the dorsal border of the prosomal segments and along the edge of the non-neurogenic ectoderm (supplementary fig. 9, Supplementary Material online).

We were able to detect expression for four of the seven identified *P. opilio* Sox genes (*SoxN*, *Sox 21b*, *SoxC*, and *SoxD*). For the three other *P. opilio* Sox genes we either failed to obtain a signal, presumably because the short sequences obtained made poor probes (*Dichaete*), or we were unable to amplify the fragment using PCR (*SoxE-1* and *SoxE2*).

*Po-SoxN* and *Pt-SoxN* are strongly expressed in the developing neuroectoderm consistent with our previous analysis of *Pt-SoxN* expression (Paese, et al. 2018a), as well as at the tips and base of the prosomal appendages (Fig. S4). *Pt-SoxC-1* and *Po-SoxC* are also expressed in the developing neuroectoderm as well as in the prosomal appendages at later stages of development, although *Pt-SoxC-1* is also expressed in a few cells where *Po-SoxC* is not detected (supplementary fig. 6, Supplementary Material online). *Po-SoxD* and *Pt-SoxD-2* also exhibit similar expression in the pre-cheliceral region, prosomal appendages and as segmental stripes (supplementary fig. 7, Supplementary Material online), but an additional domain of *Pt-SoxD-2* expression was detected in the opisthosomal appendages (supplementary fig. 7, Supplementary Material online).

Finally, we found that like *Pt-Sox21b-1, Po-Sox21b* is also expressed in a segmental pattern during early stages as well as later in the neuroectoderm (fig. 2). We were also able to detect expression for *Pt-Sox21b-2* in the last two opisthosomal segments (fig. 2) as well as a faint signal in the developing neuroectoderm that did not appear to overlap with *Pt-Sox21b-1* expression at stage 9.2 (fig. 2 and supplementary fig. 10, Supplementary Material online). These results suggest that *Sox21b* played a role in segmentation ancestrally in arachnids and that there may have been subfunctionalisation of *Sox21b* function after duplication in *P. tepidariorum*.

**Figure 2.**
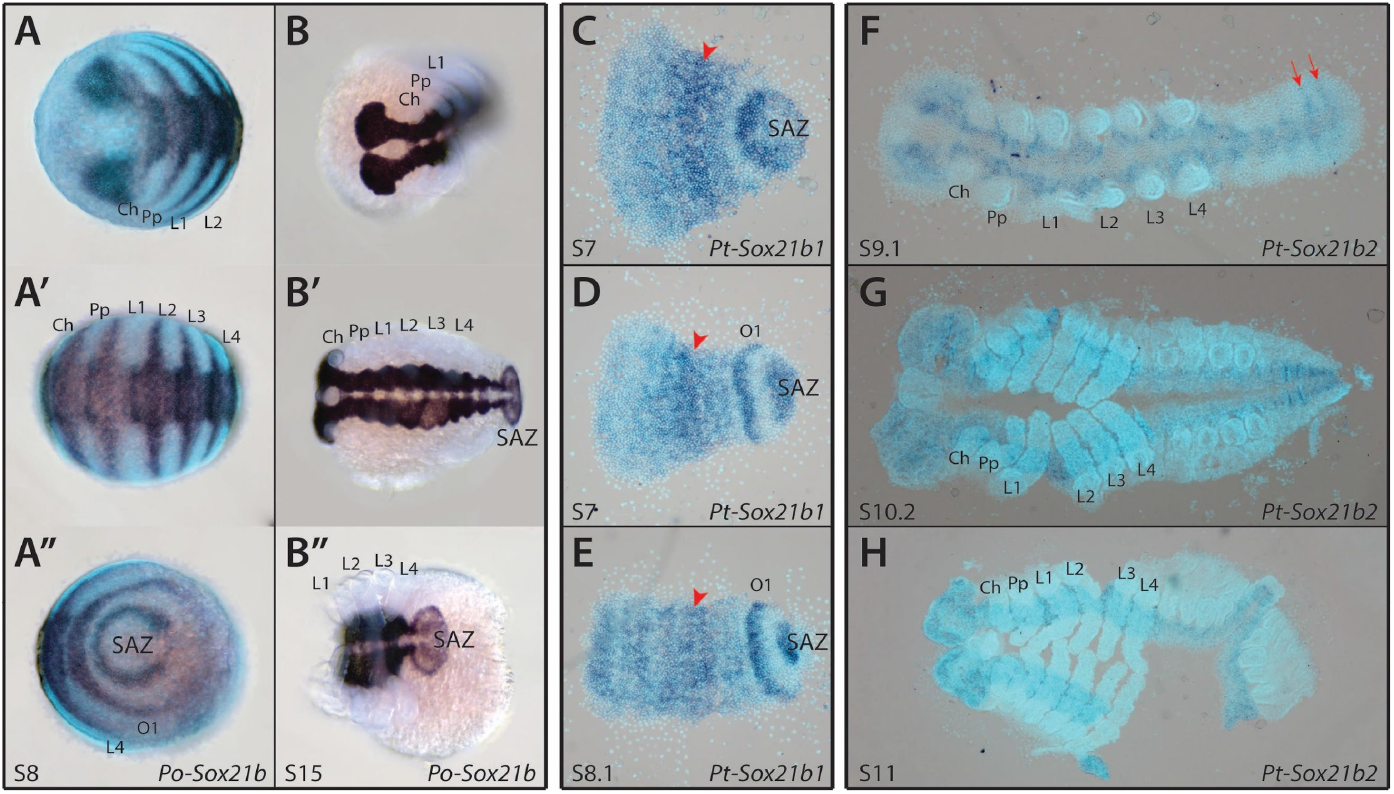
Expression patterns of *P. opilio* and *P. tepidariorum Sox21b* genes. *Po-Sox21b* has a segmental pattern of expression at stage 8 (A-A’’), similar to that previously observed for *Pt-Sox21b-1* (C-E). Expression of *Pt-Sox21b-1* in the prosomal region appears to be stronger in the presumptive L2-L4 segments (red arrowheads) at stages 7 (C and D) and 8.1 (E). Additionally, *Pt-Sox21b-1* is strongly expressed in the SAZ and first opisthosomal segment at these stages (C-E). ISH for *Pt-Sox21b-2* produced a faint signal in the developing neuroectoderm, suggesting this paralog is expressed at low levels in this tissue (F-H). Furthermore, at stage 9.1 (F), expression appears to be segmental in the last two segments in the opisthosoma (red arrows). No signal was detected for *Pt-Sox21b-2* during earlier stages of embryogenesis. Anterior is to the left in all images. Embryos in C-H are flat mounted. SAZ, segment addition zone; Ch, chelicerae; Pp, pedipalps; L1-L4, walking legs 1-4; O1, opisthosomal segment 1.

### Are other Sox genes in addition to Sox21b-1 required for segmentation in P. tepidariorum?

Our analysis of Sox gene expression suggested that in addition to *Pt-Sox21b-1*; *Pt-D, Pt-Sox21a-1* and *Pt-SoxD-2* may also be involved in segmentation in *P. tepidariorum* (supplementary figs 3, 5 and 7, Supplementary Material online). Therefore we carried out pRNAi knockdown of these three Sox genes, as well as repeating *Pt-Sox21b-1* knockdown as a positive control and for further analyses of this treatment (supplementary fig. 11 and supplementary file 3, Supplementary Material online). We did not observe any effect on embryonic development from knocking down *Pt-D, Pt-Sox21a-1* and *Pt-SoxD2,* although we were able to reproduce the same phenotypic effects at an approximately similar frequency as we previously obtained with *Pt-Sox21b-1* parental RNAi (pRNAi) (Paese, et al. 2018b) (supplementary fig. 11 and supplementary file 3, Supplementary Material online). This suggests that while *Pt-Sox21b-1* is essential for segmentation, *Pt-D, Pt-Sox21a-1* and *Pt-SoxD-2* may not be required.

### Further analysis of Sox21b-1 function in prosomal segmentation in P. tepidariorum

Knockdown of *Sox21b-1* expression in *P. tepidariorum* inhibits the formation of all leg-bearing segments (Paese, et al. 2018b). To further evaluate the effect of *Pt-Sox21b-1* knockdown on the regulation of anterior segmentation we characterised the expression patterns of *Pt-Dll*, *Pt-hb,* and *Pt-Msx1,* which have known roles in this process (Akiyama-Oda and Oda 2020; Leite, et al. 2018; Pechmann, et al. 2011; Schwager, et al. 2009).

*Pt-Dll* is required for the development of L1 and L2 segments, and is also expressed in the SAZ (Pechmann, et al. 2011) (fig. 3A-D). The early ring-like expression domain in L1 is still present in stage 5 knockdown embryos (n=5) and maintained as a stripe up to stage 7 (n=3) (fig. 3E-G). In *Pt-Sox21b-1* pRNAi embryos, at stage 8.1, the L1 stripe of *Pt-Dll* expression is somewhat weaker than in wild type embryos and the L2 stripe is no longer restricted to segmental expression (n=5) (fig. 3H-I’). This is consistent with the loss of L1 and L2 upon *Pt-Sox21b-1* knockdown and suggests this treatment prevents the splitting of *Pt-Dll* expression necessary for the formation of these segments. *Pt-Sox21b-1* knockdown also results in a reduction in *Pt-Dll* expression at the posterior of the germband at stage 7 (n=3) and subsequently this expression completely disappears (n=8), consistent with perturbed development of the SAZ in *Pt-Sox21b-1* pRNAi embryos (fig. 3G-J’).

**Figure 3.**
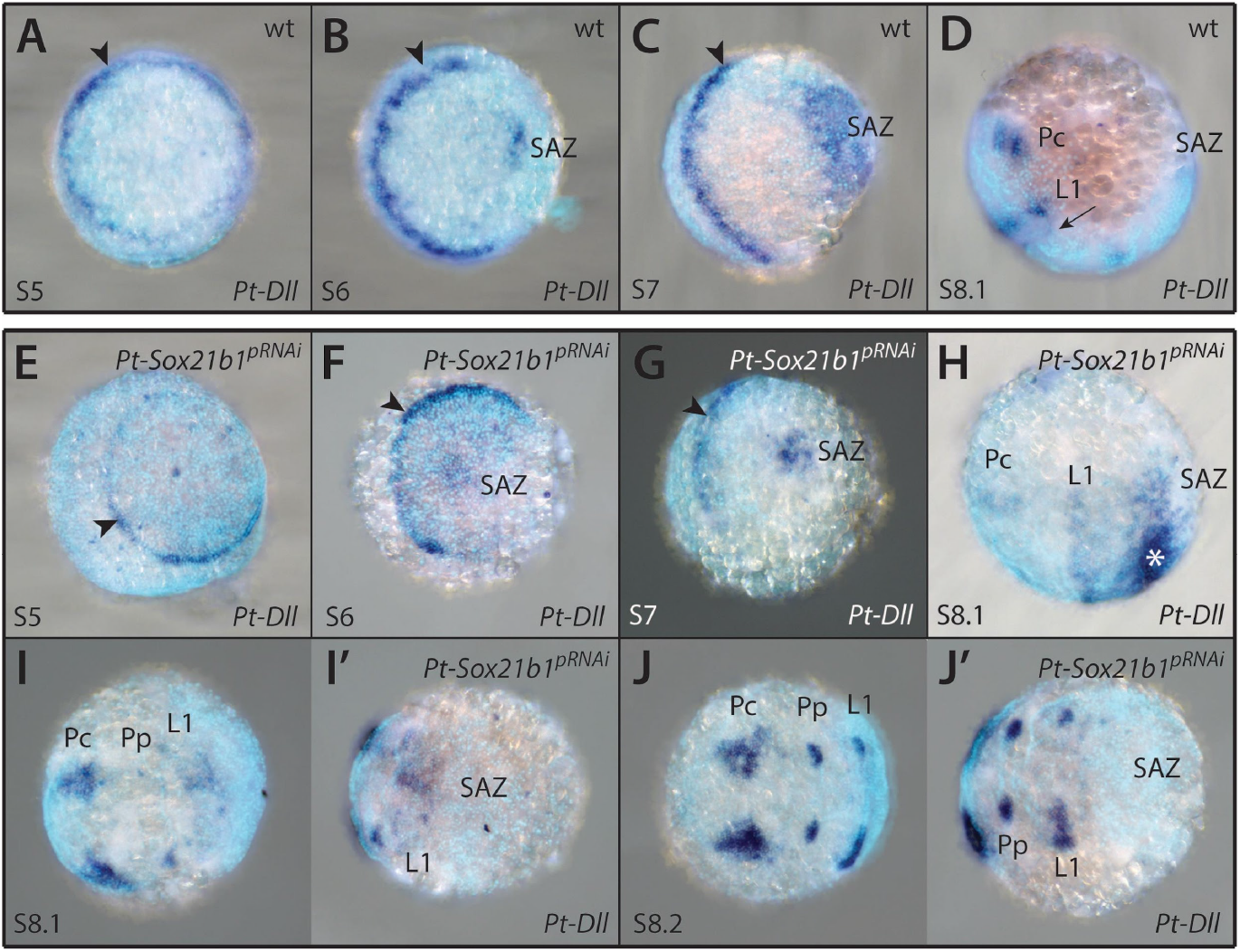
Effect of *Pt-Sox21b-1* pRNAi knockdown on *Pt-Dll* expression. *Pt-Dll* is expressed in the presumptive region of the L1 segment from as early as stage 5 (A; black arrowhead), first as a ring-like pattern which transforms into a stripe at the germ disc to germ band transition (B; black arrowhead). Additional expression domains in the SAZ and pre-cheliceral region arise at stages 6 (B) and 7 (D) respectively, as well as a faint stripe in the L2 segment (black arrow). Expression of *Pt-Dll* does not seem to be affected in stage 5 (E) and 6 (F) *Pt-Sox21b-1* pRNAi embryos. At stage 7 (G), *Pt-Sox21b-1* knockdown leads to the reduction of *Pt-Dll* expression in the SAZ, although the anterior stripe of expression is unaffected (black arrowhead). In stage 8.1 knockdown embryos (H-I’), expression is also reduced in the presumptive L1 segment and the faint stripe in the L2 segment is missing. In some embryos (n=2), ectopic expression was observed near the posterior of the germ band (H; asterisk). The pre-cheliceral and developing prosomal appendage expression domains are seemingly unaffected (H-J’). Anterior is to the left in all images. Pc, pre-cheliceral region; Pp, presumptive pedipalpal segment; L1, presumptive L1 segment; SAZ, segment addition zone.

*Pt-hb* is expressed in the prosoma (fig. 4A, B) and regulates the development of the L1, L2 and L4 segments (Schwager, et al. 2009). Several aspects of *Pt-hb* expression are disrupted by *Pt-Sox21b-1* pRNAi knockdown (fig. 4), which is consistent with the loss of segments observed after this treatment (Paese, et al. 2018b). In stage 7 *Pt-Sox21b-1* pRNAi embryos, stripes of *Pt-hb* expression corresponding to the presumptive precheliceral/pedipalpal and L1/L2 regions appear as normal, although the L4 domain appears to be perturbed in some embryos (n=7) (fig. 4C, D). Subsequently, while the pre-cheliceral domain and pedipalpal stripes of *Pt-hb* expression develop normally, the L1/L2 domain does not split into segmental stripes, and the L3 stripe cannot be distinguished from L4 (fig. 4D), while in some embryos all tissue and expression posterior of L1 is lost (n=4) (fig. 4F). These results suggest that while *Pt-Sox21b-1* is not necessary for the activation of *Pt-hb* expression, it is required for development of stripes of *Pt-hb* expression in L1 to L4 and the formation of these segments.

**Figure 4.**
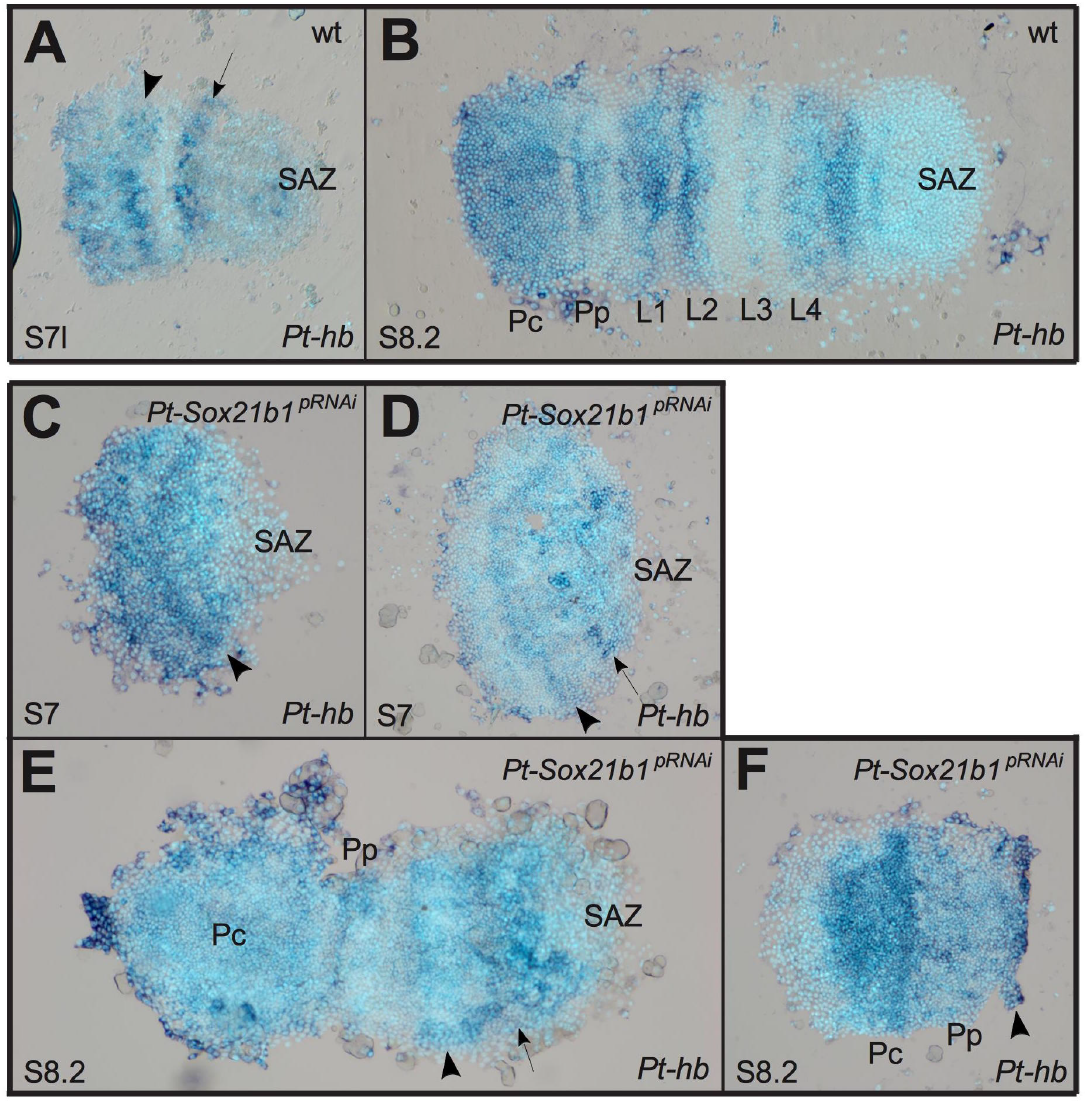
Effect of *Pt-Sox21b-1* pRNAi knockdown on *Pt-hb* expression. At late stage 7 (A), two bands corresponding to the L1/L2 segments (black arrowhead) and presumptive L4 segment (black arrow) are observed. Subsequently, by stage 8.2 (B) *Pt-hb* expression becomes segmentally restricted, with a domain covering the pre-cheliceral and cheliceral segments, and as stripes in all other anterior segments, with the strongest expression in the L1, L2 and L4 segments. Expression of *Pt-hb* in the presumptive L1/L2 segments is unaffected in stage 7 *Pt-Sox21b-1* pRNAi embryos (C and D; black arrowhead), although expression in the presumptive L4 segment is absent at times (black arrow). At stage 8.2 (E and F), expression in the pre-cheliceral and pedipalpal segments is unaffected by *Pt-Sox21b-1* knockdown, however, separation of the L1/L2 band of expression does not seem to occur (black arrowhead), and a single band of expression is observed in the presumptive L3/L4 segments (black arrow). Anterior is to the left in all images. Pc, pre-cheliceral region; Pp, presumptive pedipalpal segment; L1-L4, presumptive L1-L4 segments; SAZ, segment addition zone.

*Pt-Msx1* is expressed in a segmental pattern during early embryogenesis (fig. 5A-D) and regulates both prosomal and opisthosomal segmentation (Akiyama-Oda and Oda 2020; Leite, et al. 2018). *Pt-Msx1* is still expressed in *Pt-Sox21b-1* pRNAi embryos at stage 5, although due to the highly deformed state of the embryos analysed, it is difficult to tell whether the pattern is restricted to the presumptive L2-L4 segments or if it extends anteriorly into the presumptive head and L1 regions (n=5) (fig. 5E). In early stage 7 *Pt-Sox21b-1* pRNAi embryos, the anterior stripe in the presumptive head segments and the broad band of expression across the presumptive L2-L4 segments can still be detected (n=6) (fig. 5F-H). However, at late stage 7, while the anterior stripe of *Pt-Msx1* expression appears to be unaffected, the band of expression across the presumptive L2-L4 segments does not split into segmental stripes (n=2) (fig. 5H) (and is consistent with the effects of *Pt-Sox21b-1* embryonic RNAi (eRNAi) – see below). It was not possible to determine whether *Pt-Msx1* expression in the SAZ depends on *Pt-Sox21b-1* because the tissue posterior to L4 is lost in these embryos (fig. 5F-H). Taken together, these results are similar to the effect of *Pt-Sox21b-1* knockdown on *Pt-hb* expression and suggest that while *Pt-Sox21b-1* is not required for *Pt-Msx1* activation, it is necessary for splitting of the broad L2-L4 expression of this homeobox gene into segmental stripes in the prosoma either by directly inhibiting *Pt-Msx1* expression in some cells or perhaps indirectly by organising the prosomal cells into segments.

**Figure 5.**
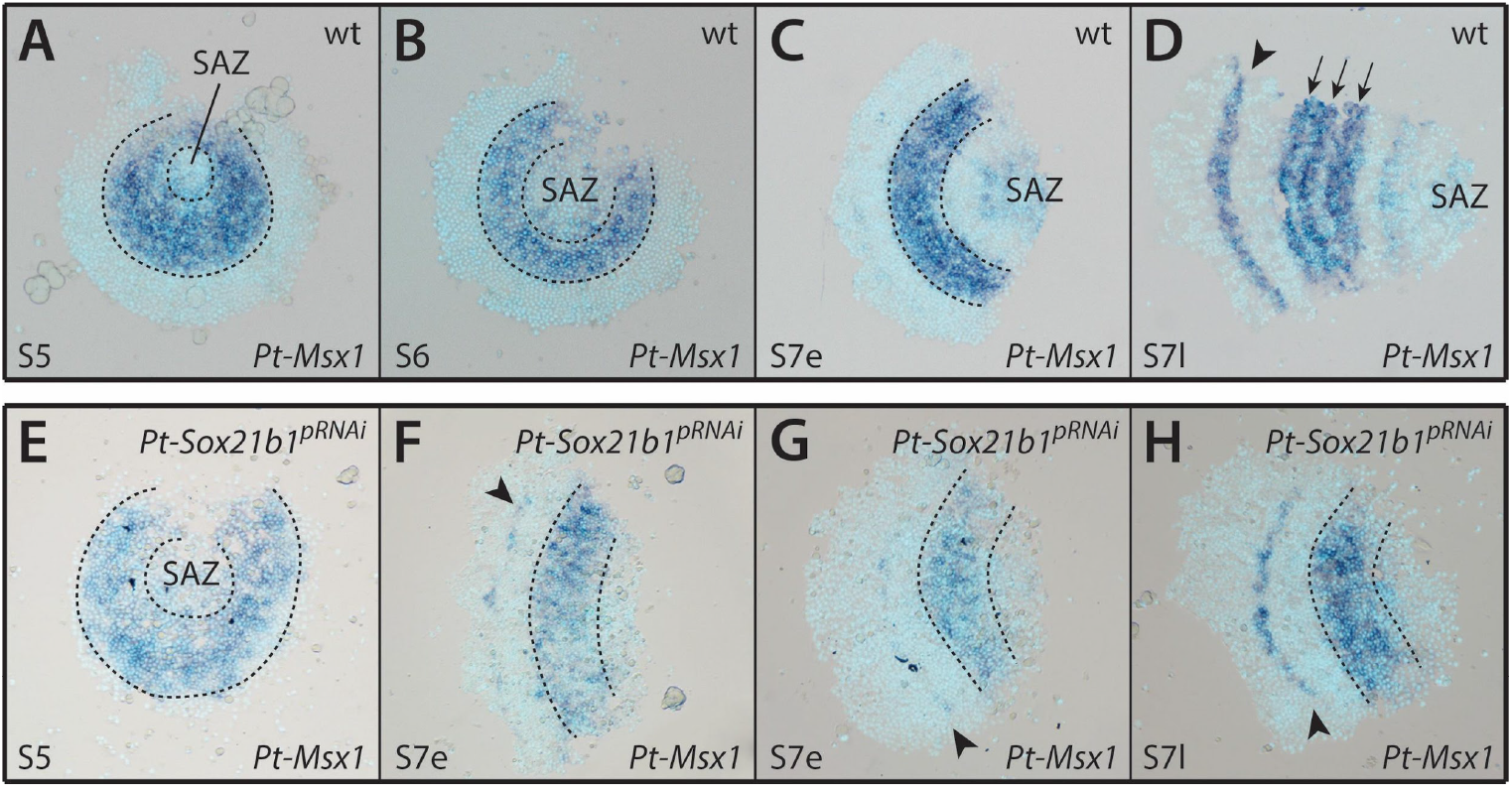
Effect of *Pt-Sox21b-1* pRNAi knockdown on *Pt-Msx1* expression. *Pt-Msx1* is first expressed at stage 5 (A) in the presumptive region of the L2-L4 segments (dashed lines), forming a band of expression at stage 6 (B). At early stage 7 (C), two additional domains of expression arise, in the SAZ and in a faint stripe corresponding to the presumptive head segments. This anterior stripe of expression becomes stronger by late stage 7 (D; black arrowhead) and the L2-L4 domain starts splitting into three stripes corresponding to each leg segment (black arrows). Faint expression is still present in the SAZ and a faint stripe can be detected in the first opisthosomal segment. Expression of *Pt-Msx1* in stage 5 *Pt-Sox21b-1* pRNAi embryos (E) is seemingly unaffected. At early stage 7 (F and G), *Pt-Sox21b-1* knockdown does not affect the band of expression in the L2-L4 presumptive region (dashed lines). However, this domain does not split into stripes in late stage 7 (H; dashed lines). The anterior stripe of expression is unaffected in stage 7 knockdown embryos (F-H; black arrowhead), although *Pt-Msx1* expression in the SAZ is lost. Anterior is to the left in all images except A and B. All embryos are flat mounted; SAZ, segment addition zone.

### Clonal analysis of Pt-Sox21b-1 knockdown in early stages of segmentation

Since pRNAi mediated knockdown of *Pt-Sox21b-1* can result in severe phenotypic effects during early embryogenesis it is sometimes difficult to ascertain direct effects of *Pt-Sox21b-1* on segmentation gene expression (Paese, et al. 2018b). To address this problem, we used eRNAi to induce *Pt-Sox21b-1* knockdown in small subsets of embryonic cells to analyse more specific and local effects of *Pt-Sox21b-1* knockdown.

We first verified the effectiveness of *Pt-Sox21b-1* eRNAi by performing ISH for *Pt-Sox21b-1* in the injected embryos (fig. 6). We observed that *Pt-Sox21b-1* expression was reduced in the cells of all clones obtained (n=12) (fig. 6D-I). In most embryos, the effect of eRNAi on *Pt-Sox21b-1* expression appeared to extend to cells adjacent to the clone region (n=11) (fig. 6D-I); a non-autonomous effect was also previously reported for eRNAi with other genes (Kanayama, et al. 2011; Schönauer, et al. 2016). Interestingly, at stages 7 (n=3) (fig. 6E) and 8.1 (n=7) (fig. 6F, G), clones in the presumptive leg segments appeared to cause a distortion of the germ band along the AP axis that extended posteriorly into the opisthosoma as a consequence of fewer cells in the affected area compared to regions adjacent to the clone. These results again highlight that the effect of *Pt-Sox21b-1* loss on cells should be considered when interpreting the effects of *Pt-Sox21b-1* knockdown on the expression of other segmentation genes.

**Figure 6.**
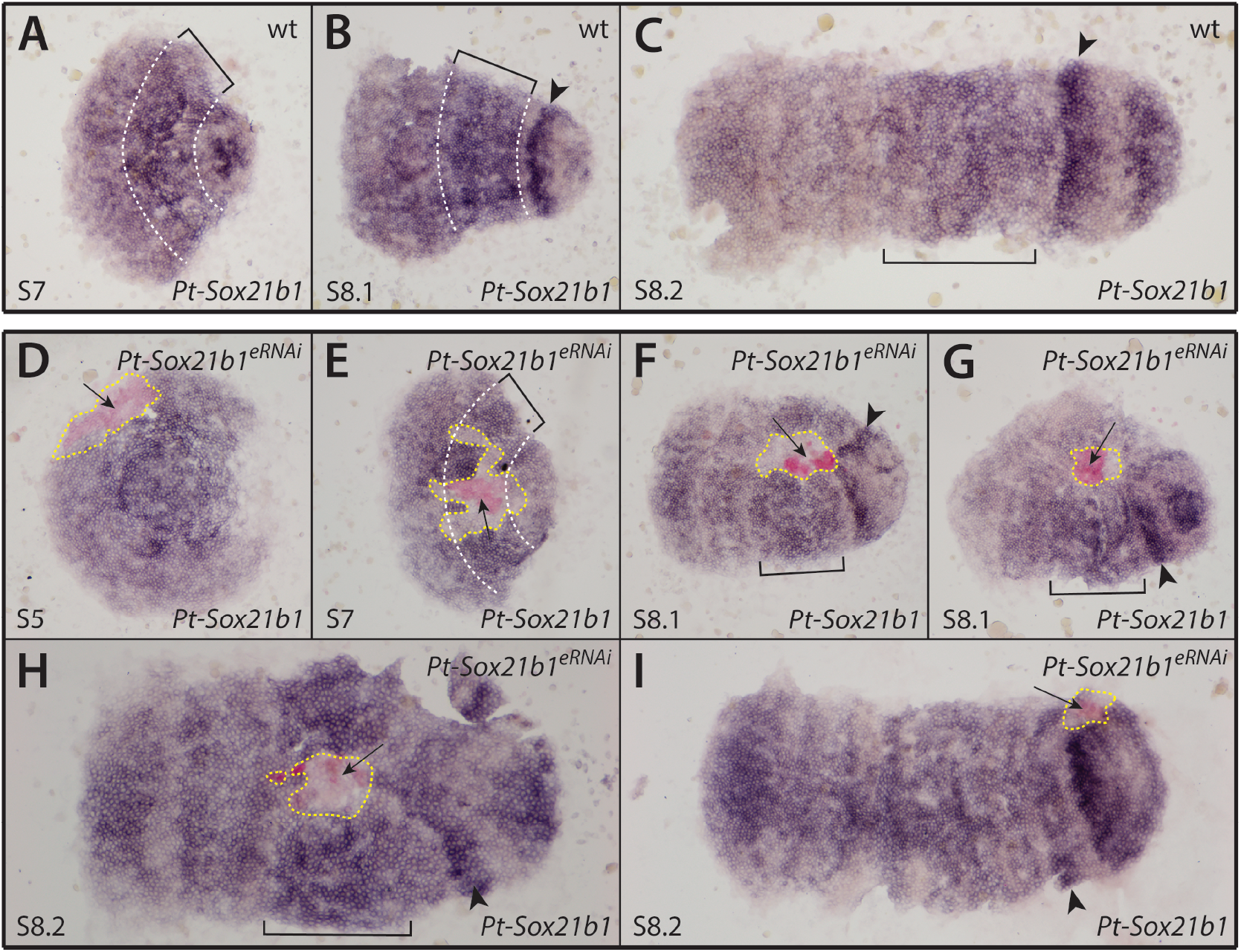
Effect of *Pt-Sox21b-1* eRNAi knockdown in early segmentation. (A-C) Expression pattern of *Pt-Sox21b-1* in stage 7 (A), 8.1 (B) and 8.2 (C) wildtype embryos. (D-I) Expression pattern of *Pt-Sox21b-1* in stage 5 (D), 7 (E), 8.1 (F, G) and 8.2 (H, I) *Pt-Sox21b-1* eRNAi embryos. Anterior is to the left in all images. All embryos are flat mounted. Dashed white lines and brackets mark the presumptive L2-L4 segments. Arrowheads: presumptive O1 segment. Arrows and yellow dashed lines indicate the knockdown clones.

### Expression of Pt-Sox21b-1 with respect to other spider segmentation genes

To better understand the regulation of prosomal and opisthosomal segmentation in *P. tepidariorum* we mapped the expression of *Pt-Sox21b-1* together with *Pt-Delta* or *Pt-caudal* via double fluorescent ISH. We chose these genes because *Pt-Dl*, like *Pt-Sox21b-1*, is expressed in the developing prosoma as well as dynamically in the opisthosoma in a similar pattern to *Pt-cad* (McGregor, et al. 2008; Oda, et al. 2007; Schönauer, et al. 2016). Moreover, although expression of both *Pt-Dl* and *Pt-cad* is lost upon *Pt-Sox21b-1* pRNAi it was unknown where and when the dynamic expression of these two key segmentation genes overlapped with that of *Pt-Sox21b-1* (Paese, et al. 2018b).

*Pt-Dl* and *Pt-Sox21b-1* are co-expressed in the prosoma at stage 7 (fig. 7A-A’’), but their expression dynamics are largely out of phase in the SAZ (fig. 7A’,A’’). At stage 8.1 *Pt-Sox21b-1* is strongly expressed in the anterior SAZ (fig. 7B’), ubiquitously in the prosoma and in a stripe domain in the forming head lobes (fig. 7B,B’). At this stage, *Pt-Dl* expression in the anterior SAZ and the cheliceral/pedipalpal region does not overlap with *Pt-Sox21b-1* expression (fig. 7B). Overall these double ISH experiments show that *Pt-Dl* and *Pt-Sox21b-1* may work together to specify prosomal segments but they are out of phase with each other in the SAZ.

**Figure 7.**
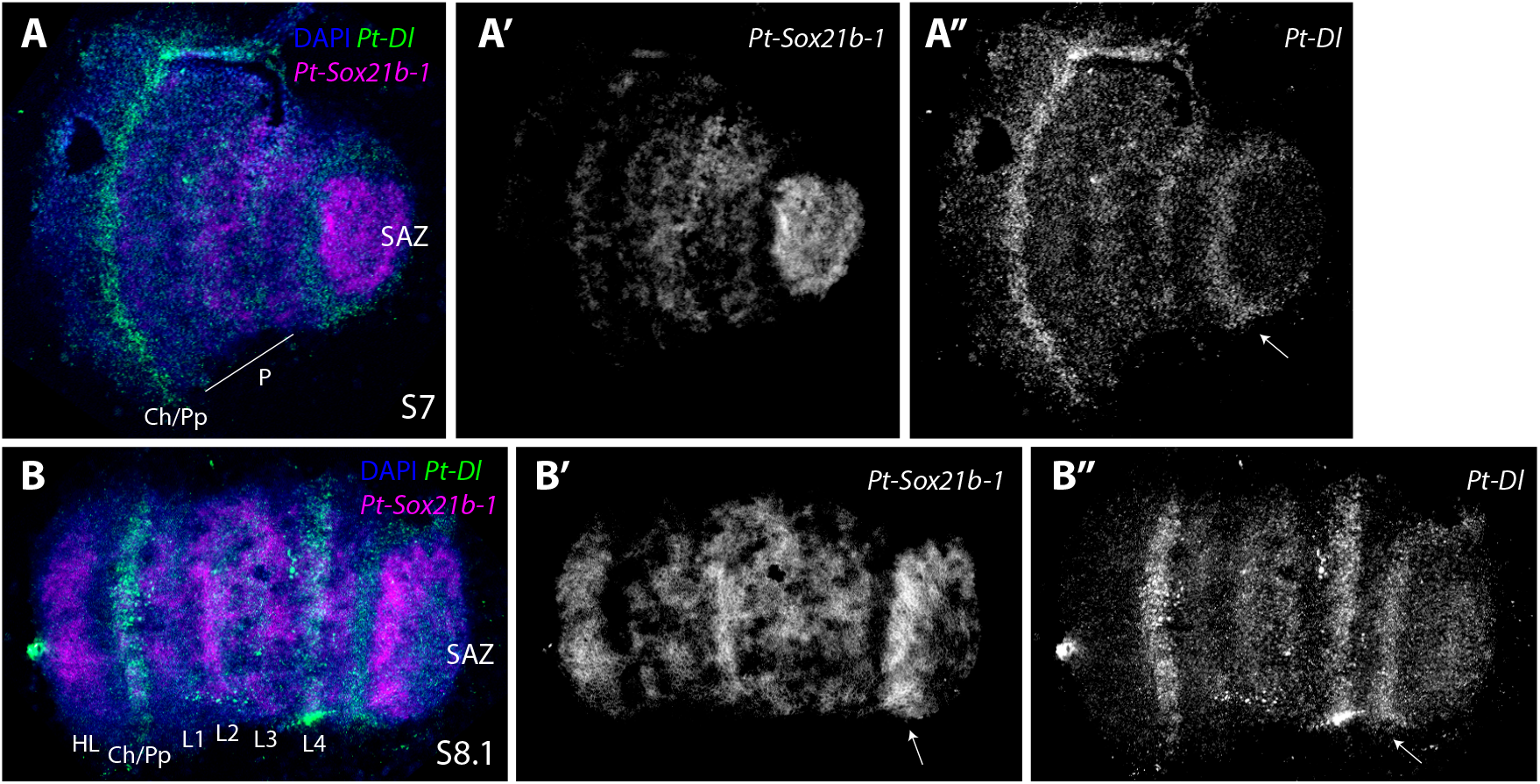
Expression of *P. tepidariorum Sox21b-1* and *Dl*. Double-fluorescent in situ of *Pt-Sox21b-1 and Pt-Dl* in stage 7 (A-A”) and 8.1 (B-B”) embryos. At stage 7 *Pt-Sox21b-1* is expressed strongly in the SAZ and throughout the forming prosoma (A’). *Pt-Dl* is also expressed in the anterior SAZ (white arrow in A’’), but this expression does not overlap with *Pt-Sox21-b1* (white arrow in A’’). *Pt-Dl* and *Pt-Sox21b-1* are co-expressed in the forming prosoma at stage 7 (A). *Pt-Dl* also exhibits a strong expression domain in the developing cheliceral-pedipalpal region (5 A’’). At stage 8.1 *Pt-Sox21b-1* is strongly expressed in the anterior SAZ (arrow in B’), ubiquitously in the prosoma and in a stripe domain in the forming head lobes (B, B’). At this stage *Pt-Dl* is expressed in the anterior SAZ (white arrow in B’’), strongly in L4 and weaker in the remaining leg-bearing segments and again strongly in the cheliceral/pedipalpal region. Anterior is to the left in all images. P, prosoma; SAZ, segment addition zone; Ch, chelicerae; Pp, pedipalps; HL, head lobe; L1-L4, walking legs 1-4.

In the case of *Pt-cad,* at late stage 6, the expression of this gene overlaps with *Pt-Sox21b-1* in the SAZ (fig. 8A’,A’’) with *Pt-cad* expression being particularly strong in the anterior region. In contrast to *Pt-Sox21b-1, Pt-cad* is not expressed in the prosoma during early stages. By stage 7 (fig. 8B-B’’) the anterior SAZ expression of *Pt-cad* appears even stronger and, as was the case with *Pt-Dl, Pt-Sox21b-*1 expression is absent from this domain. At stage 8.1 *Pt-cad* is expressed in the posterior SAZ, extending into the anterior region where it partially overlaps with *Pt-Sox21b-1* (fig. 8C,C’’). *Pt-cad* is also expressed in lateral regions of the anterior SAZ (fig. 8C’’), as well as in a lower cell layer of the fourth walking leg segment, but these domains do not overlap with *Pt-Sox21b-1* expression (fig. 8C). Taken together, the relative expression of *Pt-Sox21b-1* and *Pt-cad* suggest these genes may contribute to defining the different regions of the SAZ, perhaps working together in some posterior cells but likely playing different roles in the anterior SAZ. Furthermore, the expression patterns suggest that *Pt-Sox21b-1* regulates prosomal segment formation but the role of *Pt-cad* is restricted to sequential production of opisthosomal segments (Schönauer, et al. 2016).

**Figure 8.**
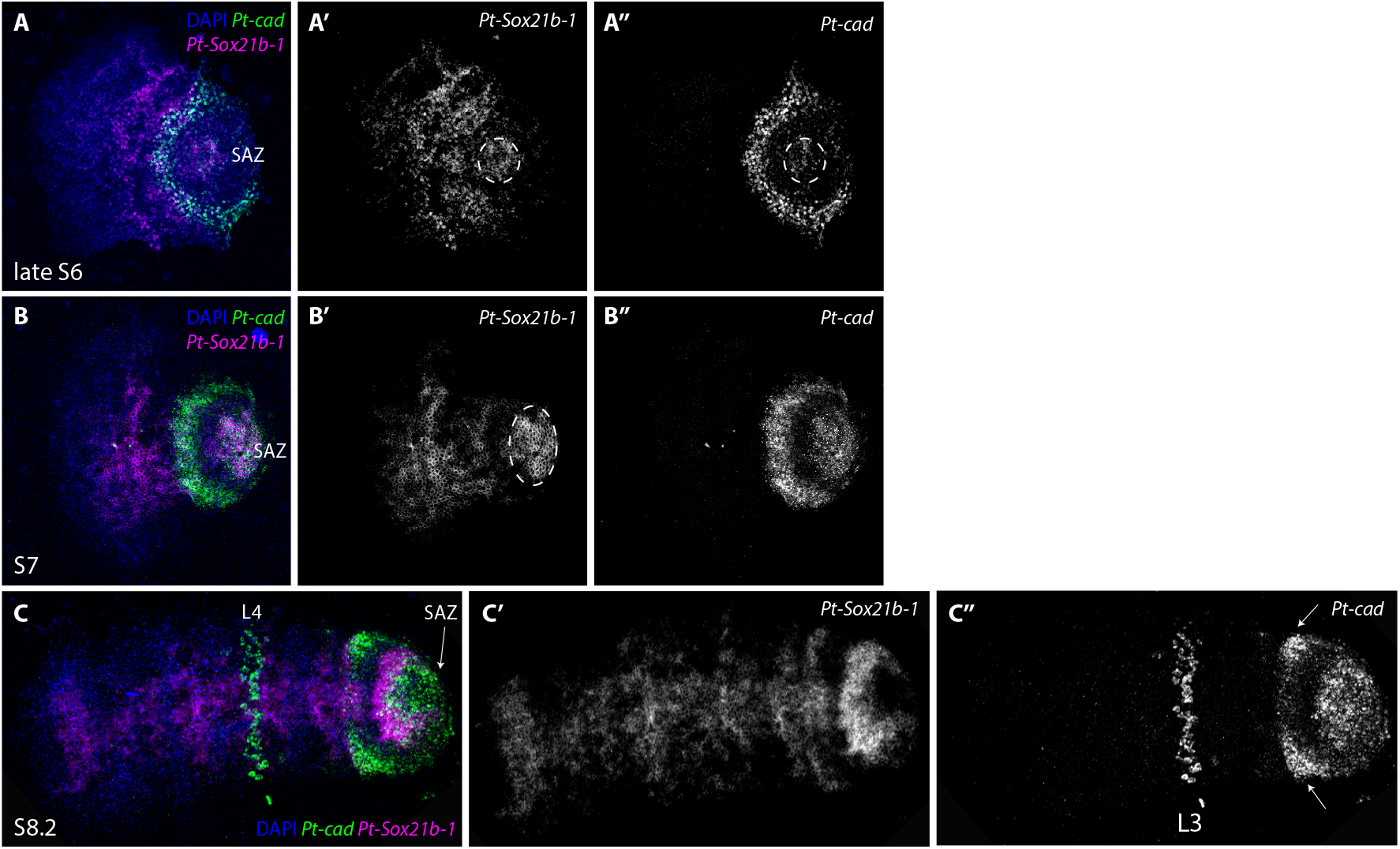
Double-fluorescent in situ of *P. tepidariorum Sox21b-1* and *cad*. Double-fluorescent in situ of *Pt-Sox21b-1 and Pt-cad* in stage 6 (A-A”), 7 (B-B”) and 8.1 (C-C”) embryos. (D-F”). At late stage 6 *Pt-Sox21b-1* is expressed in a faint circular domain in the posterior SAZ overlapping with *Pt-cad* (dashed circle in A’’), although expression of these two genes doesn’t overlap in the anterior SAZ (A, A’’). *Pt-Sox21b-1* and *Pt-cad* are both expressed in a solid domain the posterior SAZ at stage 7 (B). *Pt-cad* is not expressed in the prosoma at stage 7 (B’), but a second strong expression domain is observed in the anterior SAZ, which does not overlap with *Pt-Sox21b-1* expression (B, B’). At stage 8.2 *Pt-cad* expression can be observed in the anterior and posterior SAZ, where it partially overlaps with *Pt-Sox21b-1* (C, C’’). *Pt-cad* is also expressed in the lateral parts of the anterior SAZ (white arrows in C’’), as well as in the mesoderm of the fourth walking leg segment but these domains do not overlap with *Pt-Sox21b-1* expression (C). Anterior is to the left in all images. SAZ, segment addition zone; L4, 4^th^ walking leg.

### Clonal analysis of the effect of Pt-Sox21b-1 knockdown on Pt-Dl expression

We found that *Pt-Sox21b-1* and *Pt-Dl* overlap in their expression in the prosoma but they are out of phase in the developing opisthosoma (fig. 7). The loss of *Pt-Dl* expression during the development of both tagmata when *Pt-Sox21b-1* is knocked down using pRNAi suggests that, while prosomal expression of *Pt-Dl* is dependent on *Pt-Sox21b-1*, perhaps even directly, opisthosomal expression of *Pt-Dl* is only lost indirectly in *Pt-Sox21b-1* pRNAi embryos because the SAZ doesn’t develop properly (Paese, et al. 2018b). To test this further we carried out ISH for *Pt-Dl* in embryos with *Pt-Sox21b-1* eRNAi clones.

At stage 7, *Pt-Dl* is expressed in the presumptive cheliceral/pedipalpal segment, leg-bearing segments L2-L4 and in the SAZ (fig. 9A). At this stage, *Pt-Sox21b-1* knockdown leads to the loss of *Pt-Dl* expression in the developing L2-L4 segments (n=5) (fig. 9B-F). Furthermore, we also observed that the *Pt-Sox21b-1* knockdown again leads to distortion of the prosomal tissue, which is apparent in constricted *Pt-Dl* expression surrounding the *Pt-Sox21b-1* knockdown clone area (fig. 9B, E, F). During stage 8, *Pt-Dl* expression in *Pt-Sox21b-1* knockdown clones in the head region appeared normal (n=4) (fig. 10B-D and supplementary fig. 12, Supplementary Material online). However, we again observed loss of *Pt-Dl* expression in the L2-L4 segments, as well as distortion of the surrounding tissue at this stage (n=3) (Fig. 10B, D and supplementary fig. 12, Supplementary Material online).

**Figure 9.**
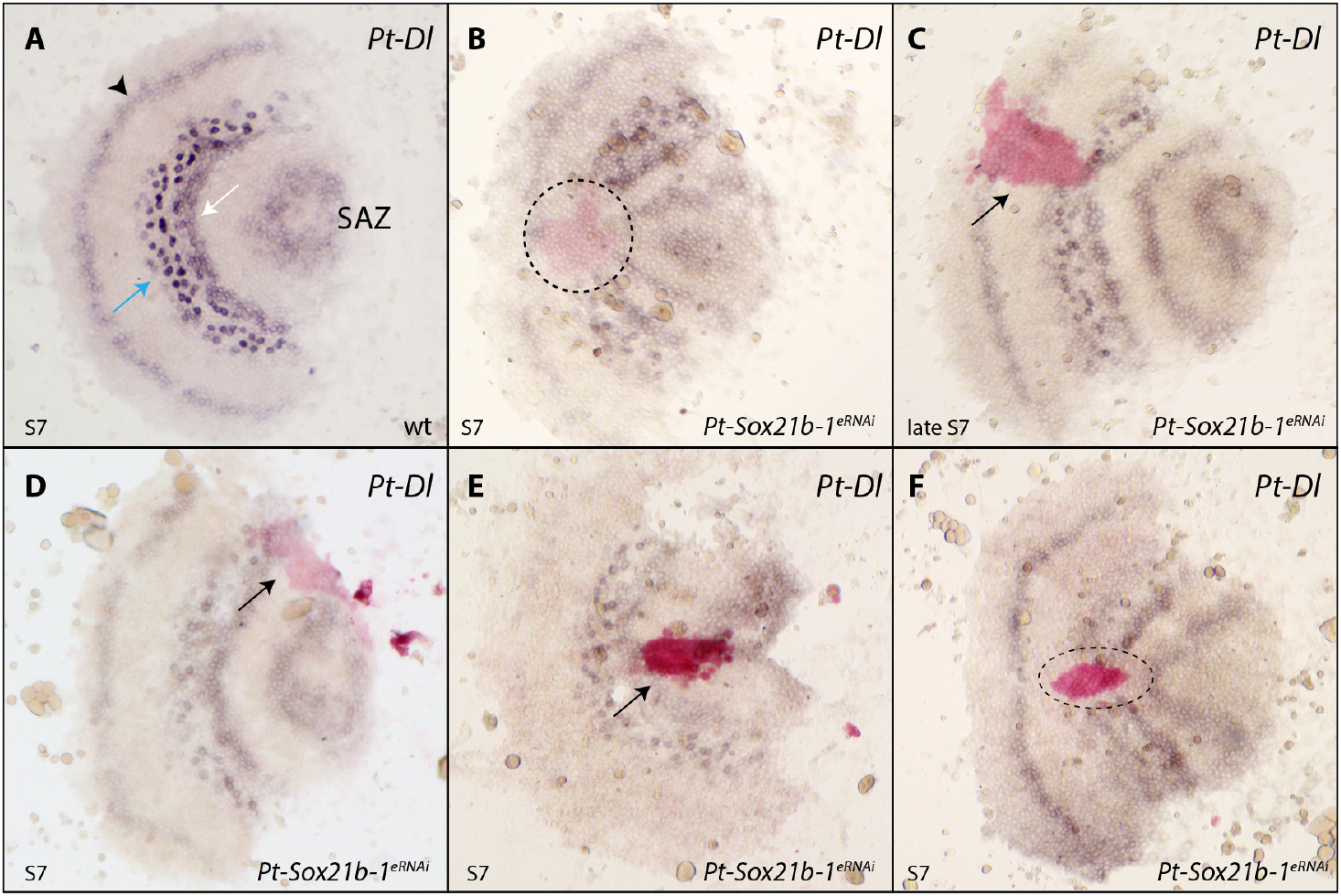
Effect of *Pt-Sox21b-1* eRNAi knockdown on *Pt-Dl* expression during early prosomal development. At stage 7 (A), *Pt-Dl* expression clears from the center of the SAZ, is expressed in two adjacent bands in the forming prosoma, a more anterior salt-and-pepper band (blue arrow) and a more posterior solid band (white arrow) and is also expressed in the presumptive pedipalpal segment (black arrowhead). *Pt-Sox21b-1* knockdown causes the loss of *Pt-Dl* expression in the prosoma (B-F), as well as the distortion of *Pt-Dl* expression domains in the tissue surrounding the *Pt-Sox21b-1* clones (B, E, F). Anterior is to the left. All embryos are flat mounted. Black arrows point at knockdown clones marked by pink staining (C, D, E) and dashed lines the estimated extent of the entire knockdown region surrounding the stained area, including potential non-autonomous affects (B, F). Pp, presumptive pedipalpal segment; L1-L4, presumptive L1-L4 segments; O1, presumptive O1 segment; O2, presumptive O2 segment; SAZ, segment addition zone.

**Figure 10.**
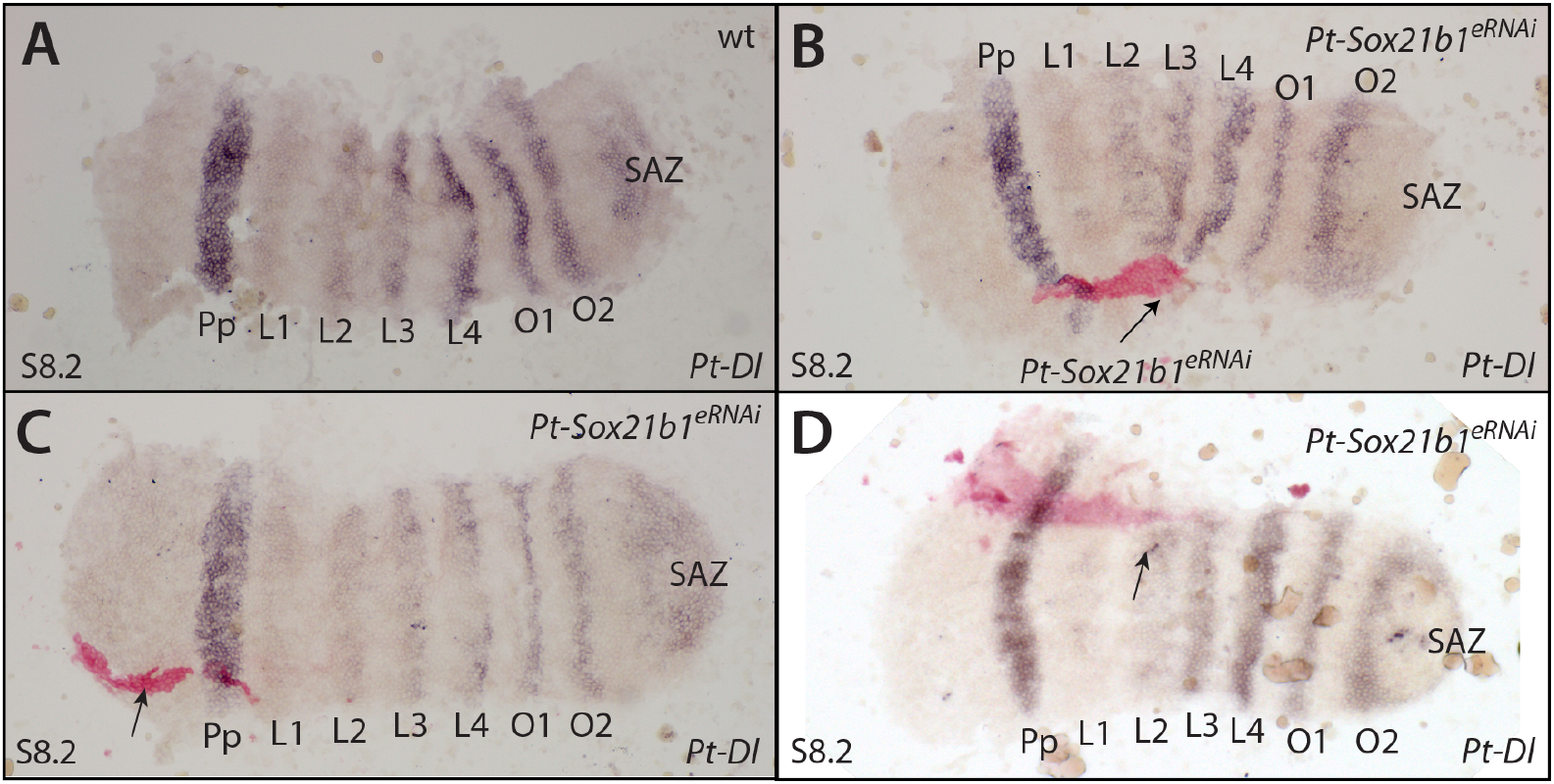
Effect of *Pt-Sox21b-1* eRNAi knockdown on *Pt-Dl* expression during later prosomal development. At stage 8.2 (A), *Pt-Dl* expression is restricted to segmental stripes of the prosoma of varying strength, the opisthosoma and the SAZ (A). *Pt-Sox21b-1* knockdown clones in the developing head do not appear to affect *P-Dl* expression (B-D). Anterior is to the left. All embryos are flat mounted. Knockdown clones are marked by pink staining and black arrows point at tissue constriction caused by *Pt-Sox21b-1* knockdown. Pp, presumptive pedipalpal segment; L1-L4, presumptive L1-L4 segments; O1, presumptive O1 segment; O2, presumptive O2 segment; SAZ, segment addition zone.

These results suggest that *Pt-Dl* expression in L2-L4 is dependent on *Pt-Sox21b-1*, and moreover, that the physical organisation of these cells into separate prosomal segments by this Sox gene is regulated via *Pt-Dl*. Unfortunately, we were unable to obtain clones in the opisthosoma but it is likely that expression of *Pt-Dl* in this tissue only indirectly requires *Pt-Sox21b-1* to regulate the formation of the SAZ.

## Discussion

### The expanded Sox repertoire of arachnopulmonates

We previously reported duplication of several Sox gene families in the spider *P. tepidariorum* and our new analysis provides a wider perspective on the evolution of the Sox genes in arachnids (Paese, et al. 2018a). We find that all arachnopulmonates we analysed have at least two copies of each Sox gene family, except for the SoxB genes *Dichaete*, *SoxN* (with the exception of *C. sculpturatus*) and *Sox21b* (with the exception of *P. tepidariorum*). This is consistent with the retention of *Sox21a, SoxC, SoxD*, and *SoxF* ohnologs after the WGD in the ancestor of arachnopulmonates that was not shared with harvestmen and ticks. Our data indicate that *SoxE* may have been duplicated in arachnids prior to the arachnopulmonate WGD because there are two copies in all arachnids surveyed including the tick and harvestman. The three copies of *SoxE* genes in *C. sculpturatus* and the two whip spider species could represent retention of a further copy after WGD or lineage specific duplications. We suggest that there may have been subsequent lineage-specific duplication of *SoxN* in *C. sculpturatus,* and *Sox21b* in *P. tepidariorum*, although we cannot completely exclude that these are true ohnologs and only one was retained in other lineages. Analysis of additional lineages may help to resolve this. Unfortunately, previous analysis of the synteny of *Pt-Sox21b-1* and *Pt-Sox21b-2* was inconclusive with respect to whether they had arisen by WGD or a more recent tandem duplication (Paese et al., 2018b). Our survey also suggests there have been lineage-specific duplications of *SoxC* and *SoxD* genes in the spiders *P. amentata* and *M. muscosa*, respectively, and perhaps other families in *C. sculpturatus* (fig. 1). This Sox gene retention in arachnids after WGD broadly parallels the retention of ohnologs of these genes after WGD in vertebrates (Schepers, et al. 2002; Voldoire, et al. 2017) and further indicates similar genomic outcomes following these independent WGD events (Leite, et al. 2018; Schwager, et al. 2017). Interestingly, the main fate of retained duplicated Sox genes in vertebrates appears to be subfunctionalisation, although there are likely cases of neofunctionalization, for example in teleosts (Cresko, et al. 2003; De Martino, et al. 2000; Klüver, et al. 2005; Voldoire, et al. 2017).

To evaluate the fates of retained spider Sox ohnologs we compared their expression during embryogenesis to their single-copy homologs in the harvestman *P. opilio*. Our comparison of Sox gene expression patterns provides evidence for broad conservation between the spider *P. tepidariorum* and the harvestman *P. opilio*, consistent with the expression and roles of these genes in other arthropods (Janssen, et al. 2018). However, differences in the expression of *P. tepidariorum* duplicates with respect to their expression in *P. opilio* could represent cases of sub- and/or neofunctionalization, but this requires verification by comparing the expression of Sox genes among other arachnids with and without an ancestral WGD. Comparison of the expression of *Sox21b* gene expression between *P. tepidariorum* and *P. opilio* indicates that these genes likely regulated segmentation in the ancestor of arachnids rather than deriving this role after WGD or tandem duplication in this spider.

### Pt-Sox21b-1 is used in different GRNs underlying prosomal and opisthosomal segmentation

*Pt-D, Pt-Sox21a-1* and *Pt-SoxD-2,* like *Pt-Sox21b-1,* are also expressed in patterns that could indicate they play a role in segmentation. Indeed it is intriguing that *Pt-D* and *Pt-Sox21a-1* are both expressed in L1 and this segment is sometimes retained after *Pt-Sox21b-1* pRNAi (Paese, et al. 2018b). However, only knockdown of *Pt-Sox21b-1* has a detectable phenotypic effect. This shows that while *Pt-Sox21b-1* is required for segmentation, it suggests that these other Sox genes may only act partially redundantly (or compensate for each other) as has been suggested for Sox genes in other animals (Heenan, et al. 2016; Phochanukul and Russell 2010; Wegner 1999). It is also possible that the pRNAi knockdown of *Pt-D, Pt-Sox21a-1* and *Pt-SoxD-2* may not have been fully penetrant. Note that the knockdown of expression of the focal gene in pRNAi experiments in *P. tepidariorum* in the absence of a phenotype is difficult to conclusively verify (e.g. see Oda, et al. 2007) and so further analysis of the function of these genes using eRNAi might be needed.

*Pt-Sox21b-1* is required for the simultaneous formation of the leg-bearing prosomal segments and the sequential production of opisthosomal segments in *P. tepidariorum* (Paese et al., 2018a) (fig. 11). Here we show that *Pt-Sox21b-1* does not appear to be necessary for the activation of *Pt-Dll*, *Pt-hb* and *Pt-Msx1*. This suggests that *Pt-Sox21b-1* could help to repress *Pt-Dll*, *Pt-hb* and *Pt-Msx1* to generate stripes from broader expression domains during prosomal segmentation and/or acts to regulate the division and viability of cells and organise them into segments (fig. 11). We suggest that these aspects of *Pt-Sox21b-1* function in the formation of the limb-bearing segments likely involves direct regulation of parts of *Pt-Dl* expression in this tagma or at least indirectly via other factors. This is consistent with the loss of prosomal segments observed following *Pt-Sox21b*-*1* pRNAi and the disruption of these segments in *Pt-Dl* pRNAi embryos (Oda et al., 2007). Intriguingly it was recently shown that *Pt-hedgehog* represses *Pt-Msx1* during segmentation but it is unclear if this role of *Pt-hh* is dependent on *Pt-Sox21b-1*, and potentially *Pt-Dl*, or not (Akiyama-Oda and Oda 2020). This shows that further work is needed to decipher how *Pt-Dl*–mediated regulation of prosomal segments is integrated with *Pt-hh*, the gap-like functions of *Pt-hb*, *Pt-Dll* and *Pt-Msx1* as well as other factors involved in cell organisation in the germband such as *Toll* genes (Akiyama-Oda and Oda 2020; Benton, et al. 2016; Pechmann, et al. 2011; Schwager, et al. 2009).

**Figure 11.**
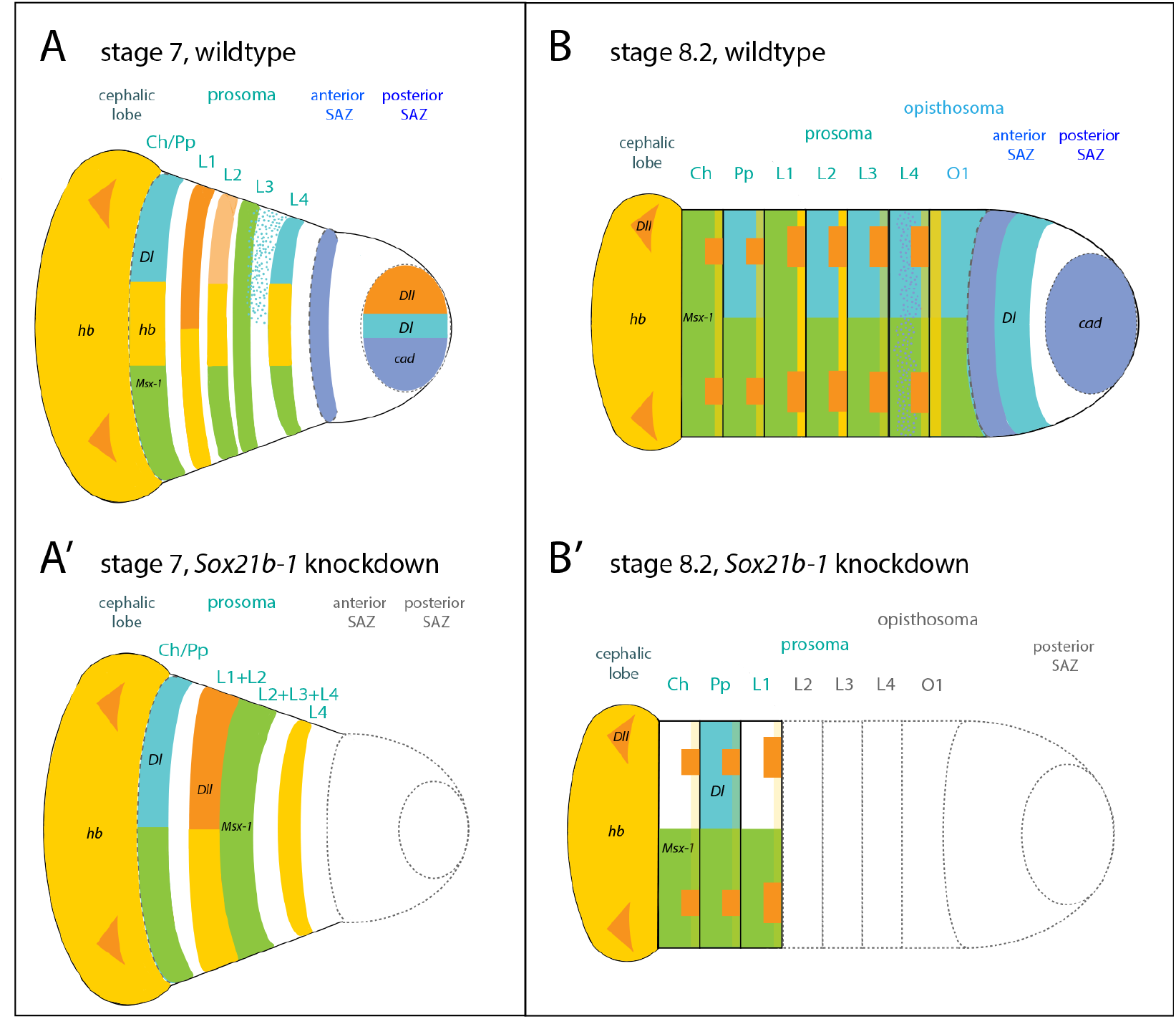
Summary of the effect of *Sox21b-1* knockdown on spider segmentation. (A,B) wildtype embryos. (A’, B’) of *Sox21b-1* knockdown embryos. At stage 7, *cad* (blue) is expressed in a circular domain in the posterior SAZ, as well as in the anterior SAZ in a stripe (A). *Dl* (turquoise) is expressed in a circular domain in the posterior SAZ, in a solid domain in L4, adjacent to an anterior salt-and-pepper domain, as well as in the cheliceral/pedipalpal segment (A). At this stage *hb* (yellow) is expressed in the prospective L1, L2 and L4, the cheliceral/pedipalpal segment and the cephalic lobe (A). *Dll* (orange) is also expressed in the posterior SAZ, L1 and in two distinct domains in the cephalic lobe (A). *Msx1* (green) is expressed in the cheliceral/pedipalpal segment, and the developing L2-L4 segments (A). In *Sox21b-1* knockdown embryos, the SAZ is missing (dashed grey lines) and expression of *Dll,* as well as *hb* expression do not split into stripes, (L1+L2) (A’). *Msx1* expression does not resolve into stripes, but presents as a solid domain (L2+L3+L4). *Dl* expression in L4 and the SAZ is lost and only remains in the cheliceral/pedipalpal segment (A’). At stage 8.2, *cad* is expressed in the posterior SAZ, in a stripe domain in the anterior SAZ and in a salt-and-pepper pattern in L4 (B). *Dl* expression can be observed in the anterior SAZ, posterior of *cad* expression and segmentally in O1, L2-L4 and the cheliceral/pedipalpal segment (B). *Msx1* is expressed segmentally in all prosomal and opisthosomal segments at stage 8.2 (Leite, et al. 2018) (B). *Dll* is expressed in the developing prosomal limb buds, and two distinct domains in the cephalic lobe (Pechmann, et al. 2011) (B). *hb* is expressed in the cephalic lobe, in all prosomal segments, whereby expression in the cheliceral, pedipalpal, L3 and L4 appear fainter, compared to expression in L1 and L2 (B). At stage 8.2, the SAZ as well as opisthosomal and prosomal segments up to L1 are lost in a *Sox21b-1* knockdown embryo (grey dashed lines) (B’). *Sox21b-1* knockdown results in down-regulation of *hb* expression in the remaining prosomal segments and only faint *hb* (light yellow) stripes remain (B’). Anterior *Msx1*, *hb, Dl* and *Dll* expression is still observed in the *Sox21b-1* knockdown embryo (B’).

*Pt-Sox21b-1* also regulates formation of the SAZ and the subsequent addition of opisthosomal segments (fig. 11). We further characterised the expression of *Pt-Sox21b-1* with respect to other genes involved in the addition of opisthosomal segments and found that *Pt-Sox-21b* expression overlaps with *Pt-cad* in some SAZ cells, but that its dynamic expression is out of phase with *Pt-Dl*. We therefore interpret the previously observed loss of *Pt-Dl* upon *Pt-Sox21b-1* pRNAi in the SAZ as a side effect of the perturbed development of this tissue rather than regulation of *Pt-Dl* by *Pt-Sox21b-1*. However, we failed to recover clones of *Pt-Sox21b-1* eRNAi knockdown in posterior cells and therefore the function and interactions of this gene in the clock-like mechanisms for sequential production of segments from the SAZ requires further investigation.

Finally, simultaneous prosomal segment production and sequential opisthosomal segment addition in spiders have similarities to the simultaneous and sequential mechanisms used in short germ and long germ insects respectively (Clark, et al. 2019). Indeed many of the same genes are involved in segmentation in spiders and insects including SoxB genes, as well as Wnt signalling, *cad* and pair-rule gene orthologs (Clark and Peel 2018; Clark, et al. 2019). However, further work is needed to better understand the gene regulatory networks underlying both prosomal and opisthosomal segmentation in spiders, including the role of *Sox21b-1* and potentially other Sox genes. It is intriguing that, while both mechanisms in spiders are regulated by *Sox21b-1*, many of the other components differ between segment formation in these two tagmata. For example *cad*, *even-skipped* and *runt* are involved in sequential but not simultaneous segmentation in *P. tepidariorum* (Schönauer, et al. 2016). Therefore while better understanding the oscillatory mechanisms used in the SAZ of spiders might add to recent new insights into understanding of how simultaneous segmentation may have evolved from sequential segmentation in insects (Clark 2017), much remains to be discovered regarding how the gap gene-like mechanism employed in the spider prosoma relates to mechanisms found insects and other arthropods.

## Materials and Methods

### Embryo collection, fixation and staging

*P. tepidariorum* embryos were collected from adult females from the laboratory culture (Goettingen strain) at Oxford Brookes University, which is kept at 25°C with a 12 hour light-dark cycle. Embryos were staged according to Mittmann and Wolff (2012) and fixed as described in Akiyama-Oda and Oda (2003). Embryos were collected and stored in RNAlater (Invitrogen) from captive mated females of the amblypygids *Charinus acosta,* at one day, one month and two months after the appearance of the egg sacs, and *Euphrynichus bacillifer,* at approximately 30% of embryonic development (Harper, et al. 2020). Mixed stage embryos were collected from a female *Pardosa amentata* (collected in Oxford) and *Marpissa muscosa* (kindly provided by Philip Steinhoff and Gabriele Uhl) and stored in RNAlater (Harper, et al. 2020). *P. opilio* embryos were collected and prepared as previously described (Sharma, et al. 2012).

### Transcriptomics

RNA was extracted from the embryos of *C. acosta, E. bacillifer, P. amentata* and *M. muscosa* using QIAzol according to the manufacturer’s guidelines (QIAzol Lysis Reagent, Qiagen) (Harper, et al. 2020). Illumina libraries were constructed using a TruSeq RNA sample preparation kit and sequenced using the Illumina NovaSeq platform (100 bp PE) by Edinburgh Genomics (https://genomics.ed.ac.uk). Raw reads quality was assessed using FastQC v0.11.9 (Andrews, 2010). Erroneous k-mers were removed using rCorrector (Song and Florea 2015) and unfixable read pairs were discarded using a custom Python script (provided by Adam Freedman, available at https://github.com/harvardinformatics/TranscriptomeAssemblyTools/blob/master/FilterUncorrectabledPEfastq.py). Reads were trimmed for adapter contamination using TrimGalore! (available at https://github.com/FelixKrueger/TrimGalore) before *de novo* transcriptome assembly with Trinity (Haas, et al. 2013). Transcriptome completeness was evaluated using BUSCO v4.0.2 (Seppey, et al. 2019) along with the arachnid database (arachnida_odb10 created on 2019-11-2; 10 species, 2934 BUSCOs) (Harper, et al. 2020). The transcriptomes will be made available on SRA upon publication.

### Identification of Sox genes

To identify the Sox gene repertoires of *C. acosta*, *E. bacillifer*, *M. muscosa*, *P. amentata*, *Centruroides sculpturatus*, *P. opilio*, *Ixodes scapularis* and *Strigamia maritima*, we performed a BLAST search (e-value 0.05) against the available genomic and transcriptomic resources (*C. acosta* – this study; *E. bacillifer* – this study; *M. muscosa* – this study; *P. amentata* – this study; *C. sculpturatus* – PRJNA422877; *P. opilio* – PRJNA236471; *I. scapularis* – PRJNA357111; *S. maritima* – PRJNA20501), using the HMG domain protein sequences previously identified in *P. tepidariorum*, *Stegodyphus mimosarum* and *D. melanogaster* (Paese, et al. 2018a). Predicted protein sequences were obtained using the ORFfinder NCBI online tool (https://www.ncbi.nlm.nih.gov/orffinder/; default settings except in ‘ORF start codon to use’ setting, where the ‘Any sense codon’ option was used to retrieve gene fragments lacking a start codon). Longest obtained ORFs were annotated as the best hits obtained from the SMART BLAST NCBI online tool (https://blast.st-va.ncbi.nlm.nih.gov/blast/smartblast/; default settings) and by reciprocal BLAST against the proteome of *P. tepidariorum*. A list of Sox sequences identified in this study and other sequences used in the subsequent phylogenetic analysis is provided in supplementary file 1, Supplementary Material online.

Sequence identity was further verified through the construction of maximum likelihood trees (supplementary figs 1 and 2, Supplementary Material online). Protein sequence alignments of HMG domains from *C. acosta*, *E. bacillifer*, *M. muscosa*, *P. amentata*, *S. mimosarum* (Paese, et al. 2018a), *P. tepidariorum* (Paese, et al. 2018a), *C. sculpturatus*, *Ph. opilio*, *I. scapularis*, *S. maritima*, *Glomeris marginata* (Janssen, et al. 2018), *T. castaneum* (Janssen, et al. 2018), *D. melanogaster* (Paese, et al. 2018a) and *Euperipatoides kanangrensis* (Janssen, et al. 2018) were generated in MEGA v.7 using the MUSCLE v.3 algorithm (default settings; Kumar, et al. (2018)). Phylogenetic analysis was performed using RAxML v.1.5b3 (Stamatakis 2014) with an LG + Γ substitution model, with nodal support inferred using the rapid bootstrapping algorithm (1000 replicates; Stamatakis, et al. (2008)). Alignments (Phylip files) and gene trees (tre files) are provided in supplementary files 2 and 4-6, Supplementary Material online. The resulting trees were visualised and processed in FigTree v1.4.4 (http://tree.bio.ed.ac.uk/software/figtree/).

### In situ hybridisation probe synthesis

Total RNA was extracted using QiAzol (Qiagen) from stage 1-14 *P. tepidariorum* embryos according to the manufacturer’s protocol. Total RNA was then used to generate cDNA using the QuantiTect reverse transcription kit (Qiagen), according to manufacturer’s guidelines. For *P. opilio*, total RNA was extracted from several clutches of stage 9-16 embryos using Trizol TRIreagent, following the manufacturer’s protocol. cDNA was generated using the Superscript III First Strand cDNA kit (ThermoFisher) with oligo-dT amplification, following the manufacturer’s protocol.

Gene-specific primers were designed using Primer3 (http://primer3.ut.ee), and T7 linker sequences were added to the 5’ end of the forward primer (GGCCGCGG) and reverse primer (CCCGGGGC). A list of primer sequences is provided in supplementary file 7, Supplementary Material online. The template for probe synthesis was generated through two rounds of standard PCR method using One*Taq*^®^ 2x Master Mix (New England Biolabs, NEB): the first PCRs from cDNA used the gene-specific primers including the T7 linker sequence. The resulting PCR product was purified using a standard PCR purification kit (NucleoSpin^®^ Gel and PCR Clean-up kit, Macherey-Nagel) and used as a template for the second PCR that used the gene-specific forward primer and a 3’ T7 universal reverse primer targeting the forward linker sequence for the antisense probe, and the gene-specific reverse primer and a 5’ T7 universal reverse primer targeting the reverse linker sequence for the sense probe. The resulting PCR products were run on an agarose gel (1-2%), and the band with the expected size excised and purified using the NucleoSpin^®^ Gel and PCR Clean-up kit (Macherey-Nagel). The second PCR products were sent for Sanger sequencing to Eurofins Genomics and checked for quality. RNA probe synthesis was performed using T7 polymerase (Roche) with either DIG RNA labelling mix (Roche) or Fluorescein RNA labelling mix (Roche), according to manufacturer’s guidelines.

### Colorimetric in situ hybridisation (ISH)

Colorimetric ISH was performed following the whole-mount protocol described in Prpic, et al. (2008) with minor modifications: steps 4-8 were replaced by two 10 minute washes in PBS-Tween-20 (0.02%) (PBS-T), and at step 18 the embryos were incubated for 30 minutes. Post-fixation was followed by ethanol treatment to decrease background: embryos were incubated for 10 minutes in inactivation buffer (75 g glycine, 600 μl 1N HCl, 50 μl 10% Tween-20 and dH_2_O to 10 ml), followed by three wash steps with PBS-T, washed 5 min in 50% ethanol in PBS-T, washed in 100% ethanol until background decreased, washed for 5 minutes in 50% ethanol in PBS-T and finally washed twice with PBS-T. Embryos were then counterstained with DAPI (1:2000; Roche) for ~20 minutes and stored in 80% glycerol in 1x PBS at 4°C. Imaging was performed using a Zeiss Axio Zoom V.16. DAPI overlays were generated in Adobe Photoshop CS6.

### Double fluorescent in situ hybridisation

Double fluorescent in situ hybridisation protocol was modified from Clark and Akam (2016): fixed embryos were gradually moved from methanol to PBS-T and washed for 15 minutes. Embryos were then transferred to hybridization buffer, hybridized overnight at 65°C and washed post-hybridization as detailed in Prpic et al. (2008). Embryos were incubated in blocking solution (Roche) for 30 minutes and AP-conjugated anti-DIG (1:2000; Roche) and POD-conjugated anti-FITC (1:2000; Roche) added and incubated for two hours. Tyramide biotin amplification (TSA Plus Biotin Kit, Perkin Elmer) was performed for 10 minutes, followed by incubation for 90 minutes in streptavidin Alexa Fluor 488 conjugate (1:500; ThermoFisher Scientific). AP signal was visualised by Fast Red staining (Kem En Tec Diagnostics). Counterstaining with DAPI (1:2000; Roche) was carried out for 5-10 minutes. Yolk granules were removed manually in PBS and germ bands were flat mounted on poly-L-lysine coated coverslips in 80% glycerol. Imaging was performed using a Zeiss LSM800 confocal. Images were processed using Adobe Photoshop CS6 and FIJI software.

### Double stranded RNA preparation

Synthesis of dsRNA was carried out using the MegaScript T7 transcription kit (Invitrogen), followed by annealing of both strands in a water bath starting at 95°C and slowly cooled down to room temperature. Purified dsRNA concentration was adjusted to 1.5-2.0 μg/μl for injections.

### Parental RNAi

Five virgin adult female spiders were injected per gene according to the protocol described in Akiyama-Oda and Oda (2006). Each spider was injected in the opisthosoma with 2 μl of dsRNA every 2 days, to a total of five injections. A male was added to the vial for mating after the second injection. Embryos from injected females were fixed at stages 5 to 8.2 for *Pt-Sox21b-1* and stages 7 - 9.2 for *Pt-Sox21a-1, Pt-SoxD-2 and Pt-D* as described above. Embryos from GFP-injected control females were generated and treated as described above. For *Pt-Sox21b-1* knockdown we used the same 549 bp dsRNA (fragment 1) as in our previous study (Paese, et al. 2018b). The same range and approximate frequencies of three phenotypic classes were observed for all cocoons from injected females (Paese, et al. 2018b). In class I embryos, all segments posterior to the first leg (L1) segment do not form properly. In class II embryos, only the head, cheliceral and pedipalpal segments are formed, and the L1 segment is missing in addition to the missing segments of class I embryos. Class III embryos do form a germ band, forming instead a disorganised cellular mass in the centre of the germ disc. Since phenotypic class III embryos do not transition from radial to axial symmetry and display major developmental defects, expression analysis was only carried out on embryos of phenotypic classes I and II (Paese, et al. 2018b). For *Sox21a-1*, *SoxD-2* and *Dichaete* parental RNAi, dsRNAs of 441 bp, 785 bp and 675 bp, respectively, were used (supplementary file 7, Supplementary Material online).

### Embryonic RNAi

Embryonic injections were carried out as described in Schönauer, et al. (2016) with minor changes. Embryos were injected between the 8- and 16-cell stages with small quantities of injection mix made up of 5 μl of FITC-dextran, 5 μl of biotin-dextran and 2.5 μl of dsRNA. Embryos were subsequently fixed at stages 5 to 8.2 of development. Visualization of eRNAi clones was achieved by detecting the co-injected biotin-dextran with the Vectastain ABC-AP kit (Vector Laboratories) after ISH, according to the manufacturer’s protocol.

## Data availability

The data underlying this article will be deposited in SRA upon publication or are already available in its supplementary material.

## Author contributions

LBG, AS, SR, LSR, PPS and APM designed this project; LBG, AS, AH, GB, MS, SA, and LSR performed the experimental work; LBG, AS, SR, AH, SA, LSR, PPS and APM analysed data; and LBG, AS and APM wrote the paper with the help of all other authors.

## Acknowledgements

This study was funded by a Leverhulme Trust grant (RPG-2016-234) to APM and AS, a NERC grant to APM and LSR (NE/T006854/1), a Nigel Groome Studentship from Oxford Brookes University to LBG, a BBSRC DTP studentship to AH, a John Fell Fund grant (0005632) from the University of Oxford to LSR and a NSF CAREER IOS-1552610 to PPS. We thank Philip Steinhoff and Gabriele Uhl for kindly providing *M. muscosa* embryos.

