## Supplementary Figures for "The evolution of Sox gene repertoires and regulation of segmentation in arachnids"

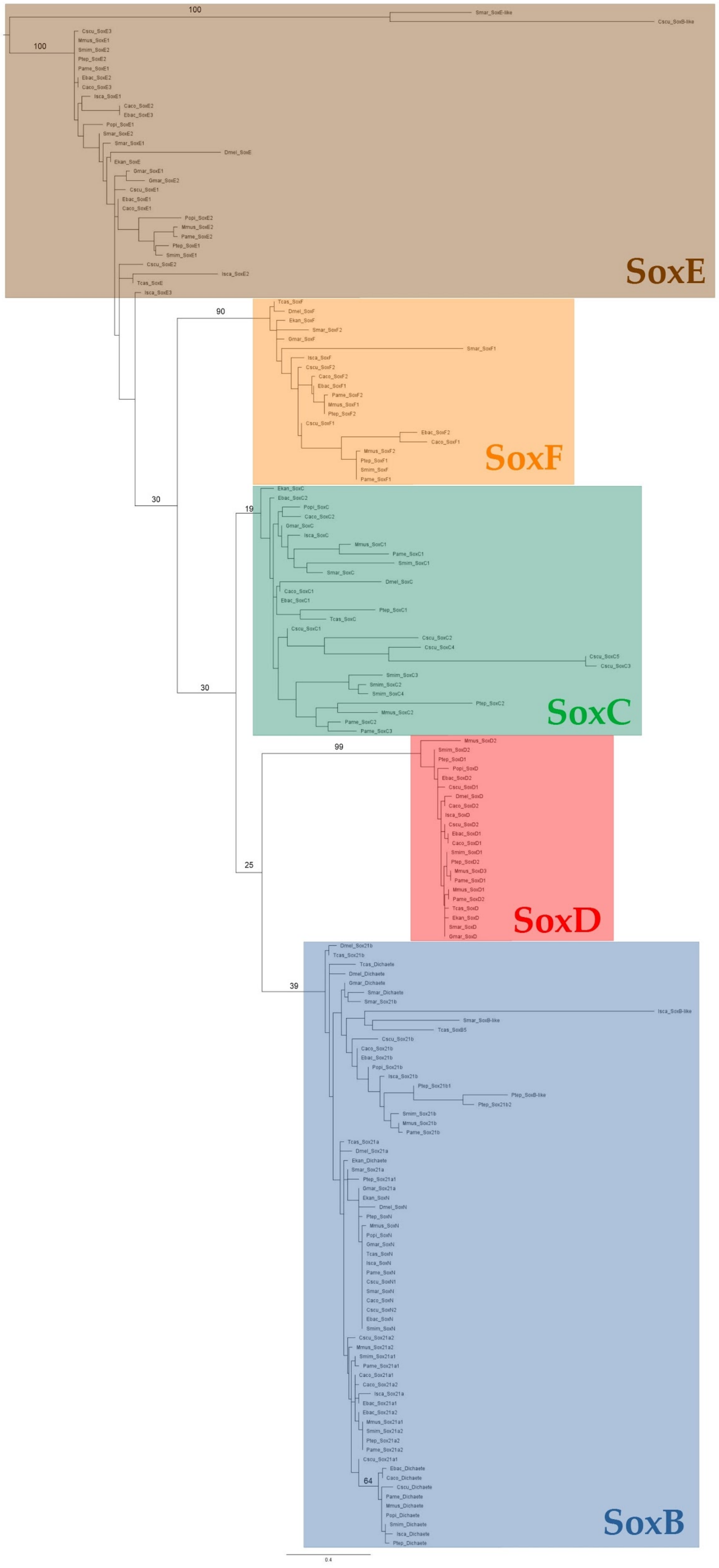


### Figure S1: Panarthropod Sox HMG domain phylogeny

Maximum likelihood tree of HMG domain sequences from *E. kanangrensis*, *G. marginata*, *S. maritima*, *T. castaneum*, *D. melanogaster*, *I. scapularis*, *Ph. opilio*, *Ce. sculpturatus*, *St. mimosarum*, *P. tepidariorum*, *M. muscosa*, *Pa. amentata*, *C. acosta* and *E. bacillifer*. Rooted at midpoint. Bootstrap values are shown for main branches. RAxML run using an LG + Γ substitution model (1000 replicates).

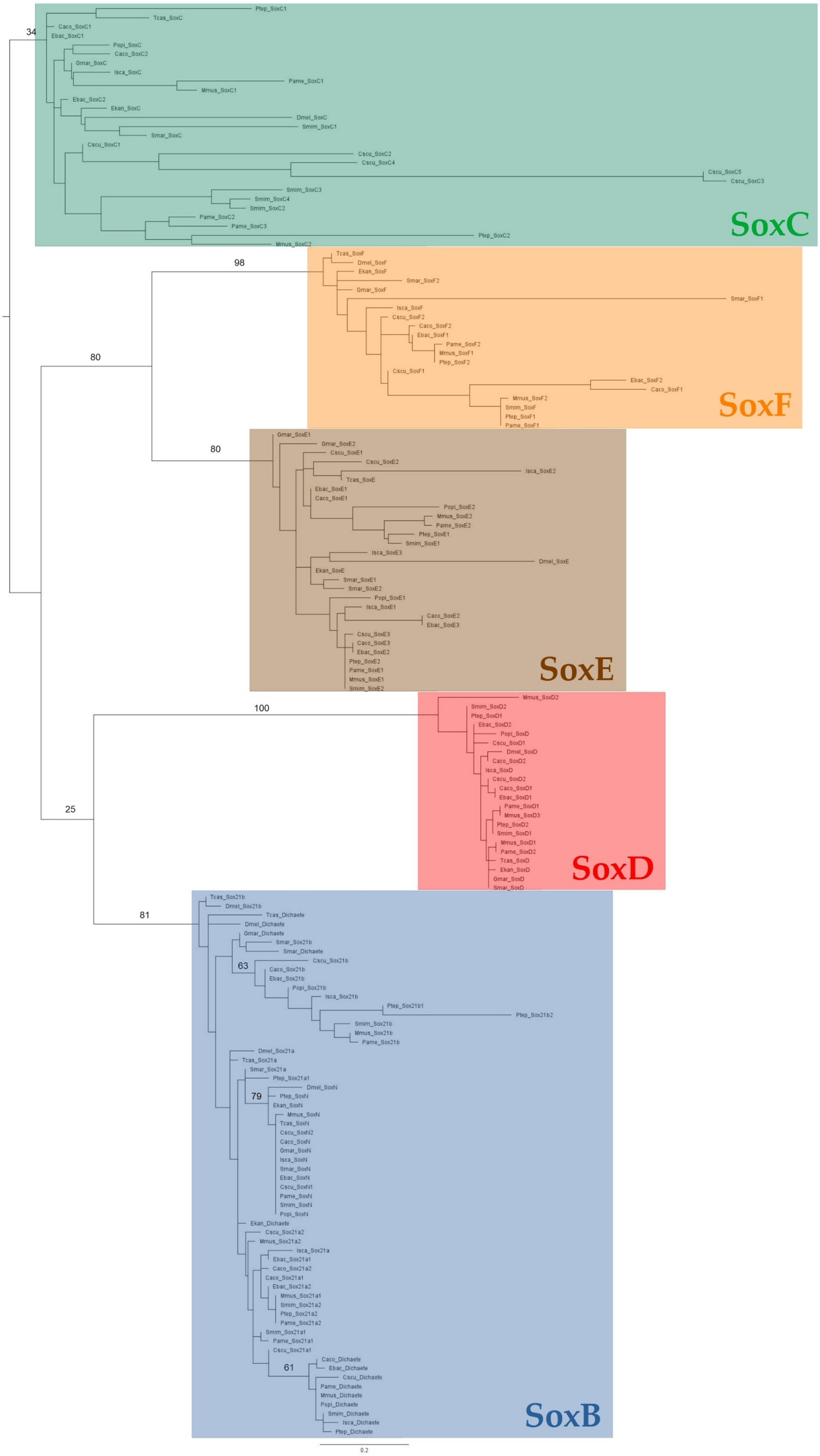


**Figure S2: Panarthropod Sox HMG domain phylogeny excluding highly divergent sequences** Maximum likelihood tree of HMG domain sequences excluding all SoxB-like and SoxElike sequences, and *Gmar-Sox21a*. Rooted at midpoint. Bootstrap values are shown for main branches. RAxML run using an LG + Γ substitution model (1000 replicates).

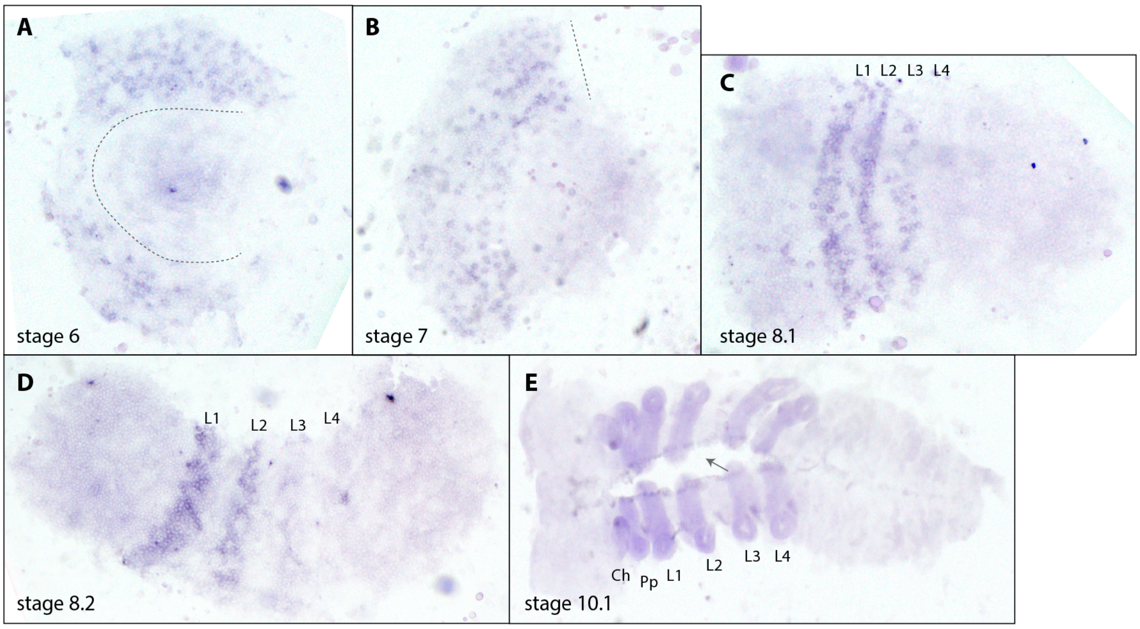


**Figure S3: Expression patterns of *P. tepidariorum Dichaete***

*Pt-D* appears to be expressed in the prosoma during embryogenesis (A-E). At stages 6 and 7, *Pt-D* is expressed in a salt-and-pepper patter throughout the developing prosoma (domain anterior of dashed lines in A; dashed line in B). At stage 8.1 *Pt-D* expression is limited to the leg bearing segments L1-L3 (C), which becomes broader at stage 8.2 (D). Later, at stage 10.1 *Pt-D* is expressed in all prosomal appendages and along the dorsal midline (arrow, E). Anterior is to the left in all images. All embryos are flat mounted. Ch, chelicerae; Pp, pedipalps; L1-L4, walking legs 1-4; O1-O5, opisthosomal segments 1-5.

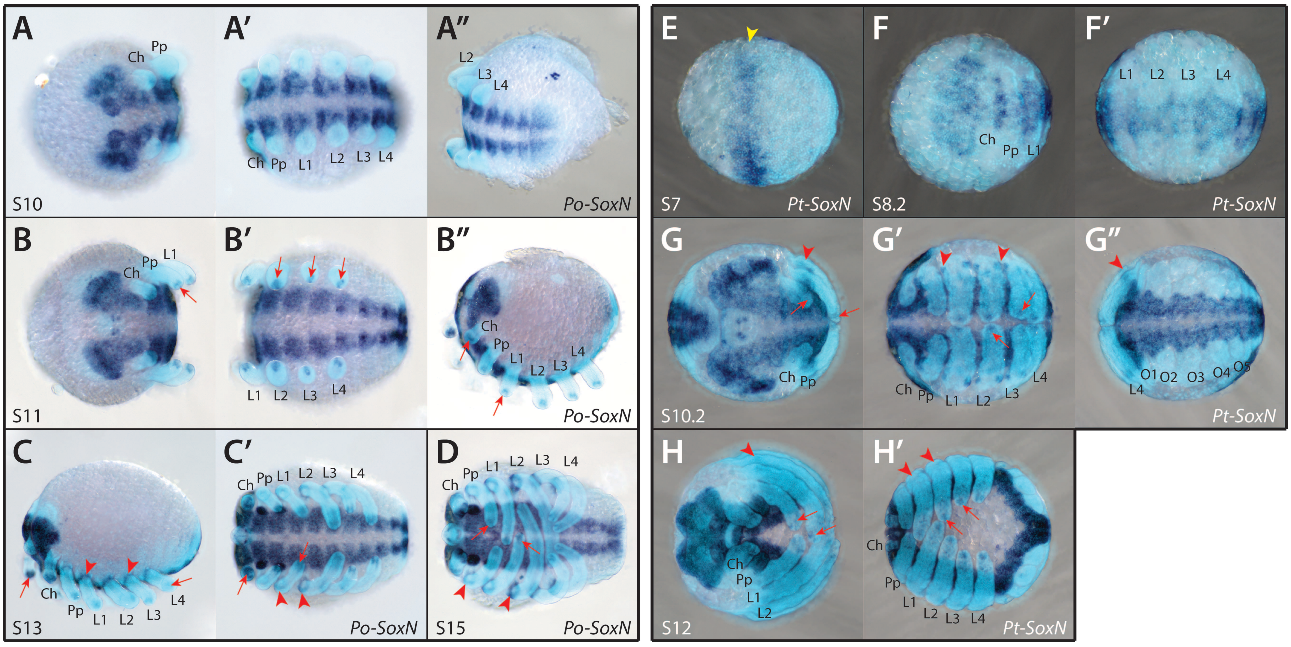


**Figure S4: Expression patterns of *P. opilio* and *P. tepidariorum SoxN* genes**

*Po-SoxN* (A-D) and *Pt-SoxN* (E-H') are both strongly expressed in the neuroectoderm throughout embryogenesis, an expression domain detected as early as stage 7 for *P. tepidariorum* in the presumptive region of the developing neuroectoderm (E, yellow arrowhead). Additional expression at the tips (red arrows) and base (red arrowheads) of each prosomal appendage is also present in both species (B-D and G-H'). Anterior is to the left in all images. Ch, chelicerae; Pp, pedipalps; L1-L4, walking legs 1-4; O1-O5, opisthosomal segments 1-5.

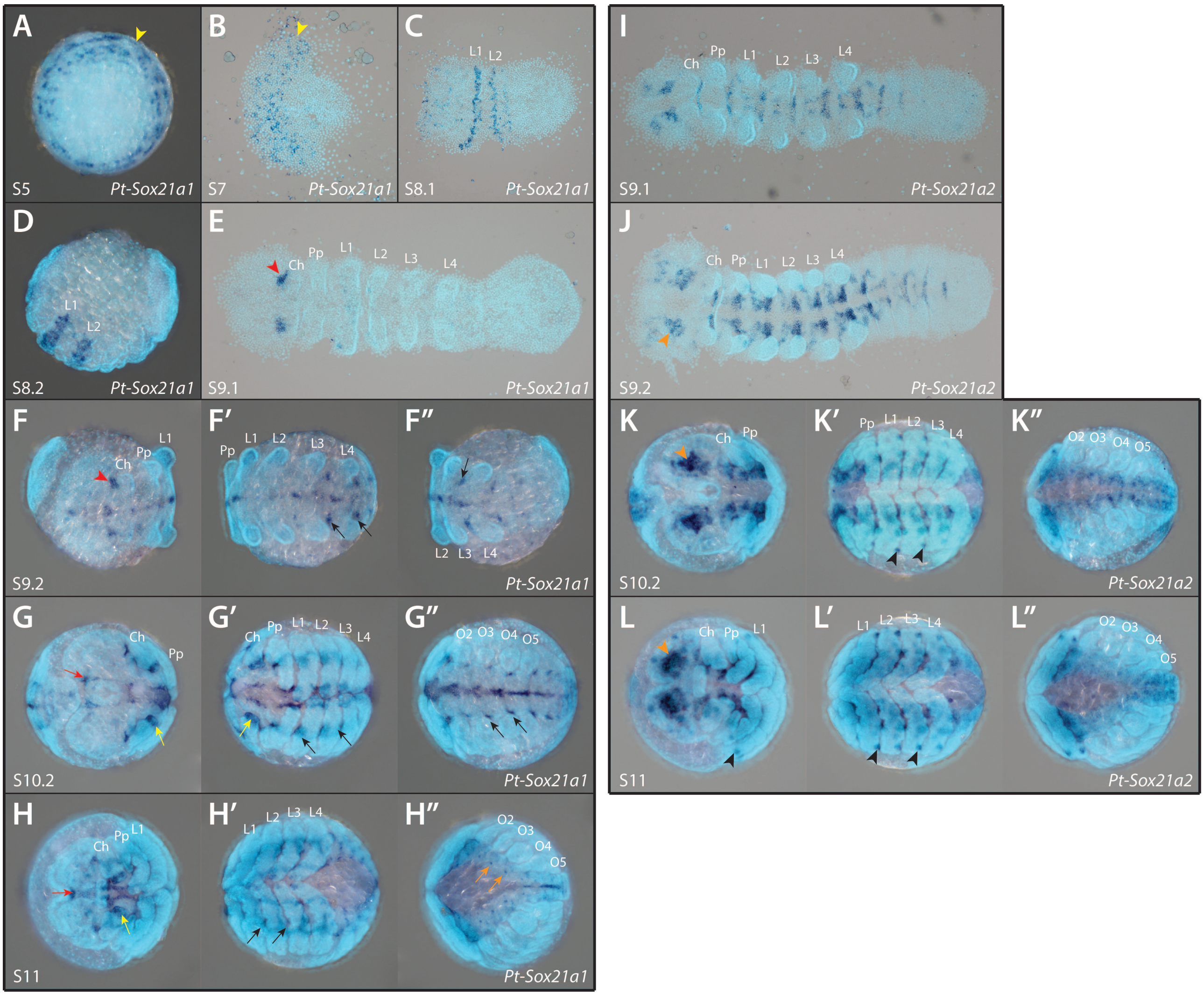


**Figure S5. Expression patterns of *P. tepidariorum Sox21a* genes**

*Pt-Sox21a-1* expression can first be observed at stage 5 (A) as a salt-and-pepper pattern around the edge of the germ disk (yellow arrowhead), which is retained after the transition from radial to axial symmetry (B). At stages 8.1 and 8.2 (C and D), expression is restricted to two bands in the L1 and L2 segments. This pattern fades by stage 9.1 (E) and a new domain of expression arises in the pre-cheliceral region (red arrowhead). At stage 9.2 (F-F''), *Pt-Sox21a-1* starts to be expressed at the midline and small clusters of cells along the dorsal edge of the developing neuroectoderm (black arrows). Expression becomes stronger by stage 10.2 (G-G''), with enlarged domains in the prosomal segments (black arrows). An additional domain can be seen at the tips of the chelicerae (yellow arrow) and surrounding the stomodaeum (red arrow). By stage 11 (H-H''), expression in the pre-cheliceral and prosomal regions increases in complexity and midline expression fades away in most of the opisthosomal segments, replaced by a dot-like pattern (orange arrows). *Pt-Sox21a-2* is mainly expressed in the neuroectoderm, starting at stage 9.1 (I) in the pre-cheliceral region and in segmental clusters of cells. This pattern is maintained throughout embryogenesis (J-L''), with increasing complexity in the developing brain (orange arrowheads) and, at later stages, an additional domain of expression is found at the base of the prosomal appendages (black arrowheads). The embryos in B, C, E, I and J were flat mounted. Ch, chelicerae; Pp, pedipalps; L1-L4, walking legs 1-4; O1-O5, opisthosomal segments 1-5.

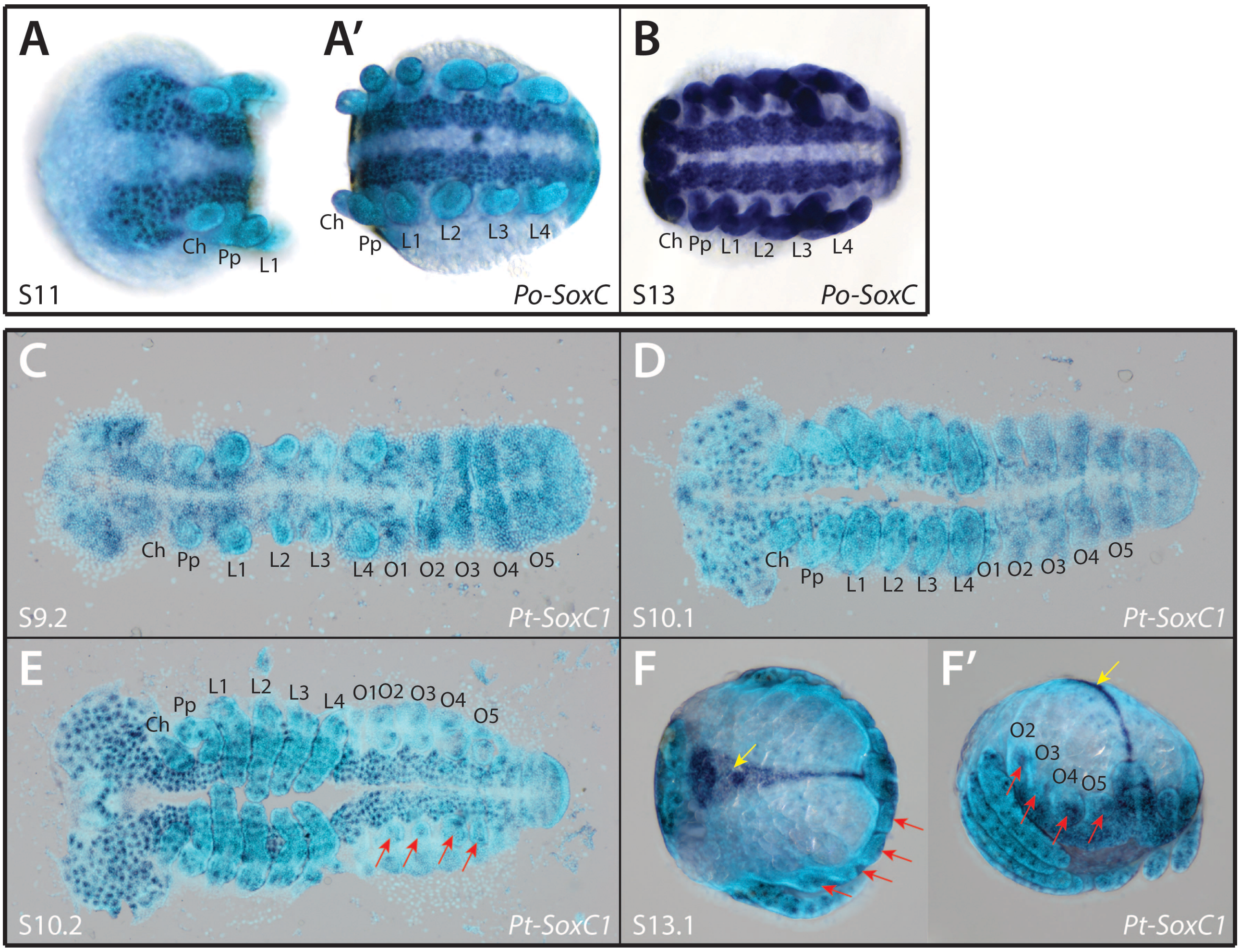


**Figure S6. Expression patterns of *P. opilio* and *P. tepidariorum SoxC* genes**

*Po-SoxC* is strongly expressed in the developing neuroectoderm, with more pronounced expression in the clusters of differentiating neuronal progenitors (A-B). At stage 13 (B), *Po-SoxC* expression was also detected in all prosomal appendages. The same neuroectodermal and prosomal appendage expression patterns were detected for *Pt-SoxC1* (C-E). Additional domains of *Pt-SoxC1* expression were observed in the opisthosomal limb buds (A-F', red arrows) and in the remaining extraembryonic tissue during dorsal closure (F and F', yellow arrows). Anterior is to the left in all images. Embryos in C, D and E were flat mounted. Ch, chelicerae; Pp, pedipalps; L1-L4, walking legs 1-4; O1-5, opisthosomal segment 1-5.

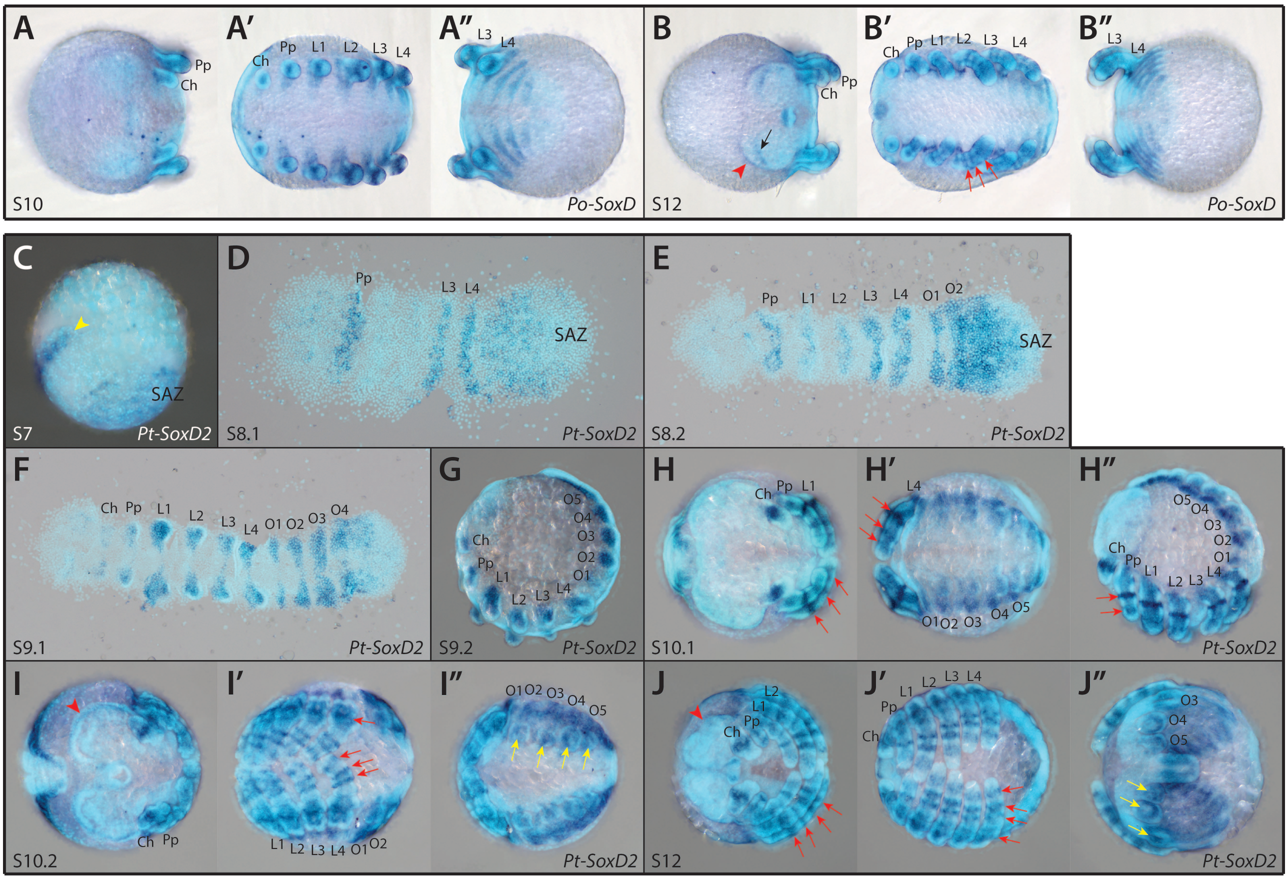


**Figure S7. Expression patterns of *P. opilio* and *P. tepidariorum SoxD* genes**

*Po-SoxD* expression is at first detected as dorsally restricted segmental stripes and in the developing prosomal appendages, forming a bifurcated pattern at the tips (A-A"). This pattern is maintained in the chelicerae at stage 12 (B-B''), developing into 2-4 rings of expression in the pedipalps and walking legs (red arrows). Expression at the edge (red arrowhead) and above the anterior furrows (black arrow) of the pre-cheliceral region is also observed at stage 12 (B), and the segmental stripe expression pattern is still present in the opisthosomal region (B''). *Pt-SoxD2* expression starts at stage 7 (C) as an anterior stripe (yellow arrowhead) and in the SAZ. At stage 8.1 (D), expression is restricted to the pedipalp, L3 and L4 segments, and anteriorly to the SAZ. Stripes of expression are continuously added to opisthosomal segments and additional stripes are then detected in the L1 and L2 segments (E and F). From stage 9.2 onwards (G-J''), *Pt-SoxD2* expression is very similar to that of *Po-SoxD*. *Pt-SoxD2* expression develops into 2-4 rings of expression in the pedipalps and walking legs (red arrows) and is also present at the edge of the pre-cheliceral region (red arrowhead), but not above the anterior furrows. Expression in the opisthosoma becomes restricted to the opisthosomal appendages (yellow arrows) and the tissue dorsal to these. Anterior is to the left in all images. The embryos in D, E and F were flat mounted. SAZ, segment addition zone; Ch, chelicerae; Pp, pedipalps; L1-L4, walking legs 1-4; O1-5, opisthosomal segment 1-5.

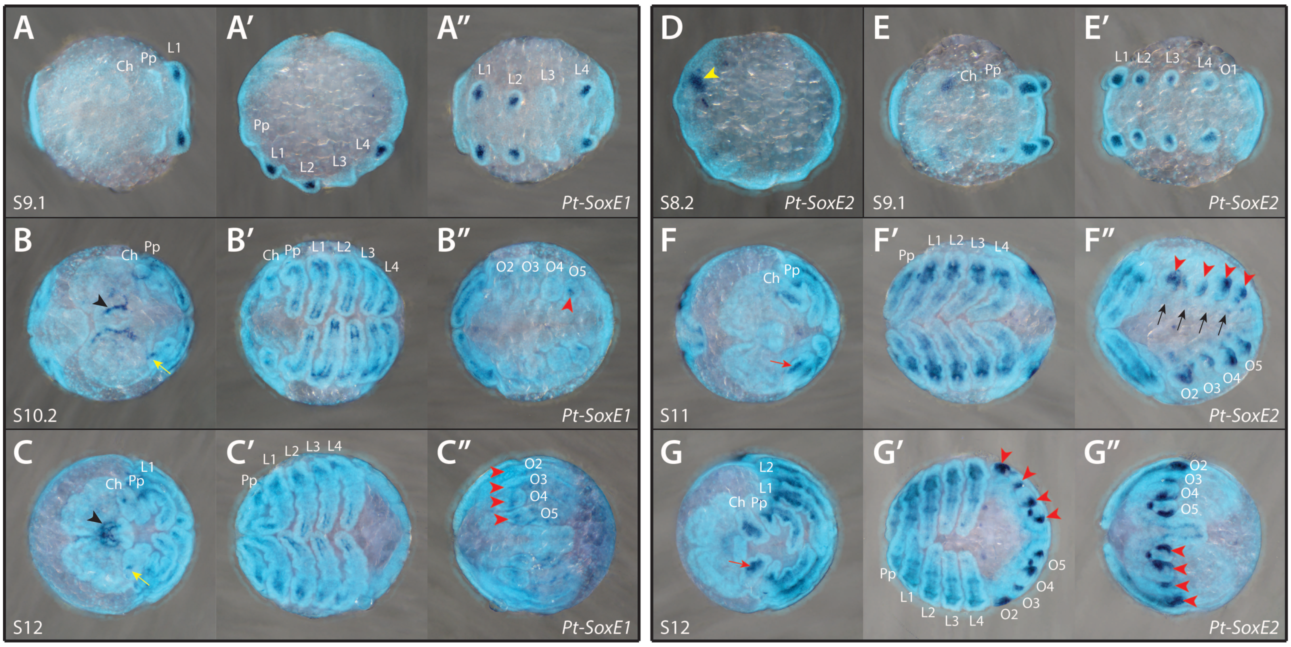


**Figure S8. Expression patterns of *P. tepidariorum SoxE* genes**

*Pt-SoxE1* expression is first visible at stage 9.1 (A-A'') in the developing prosomal appendages of the L1, L2 and L4 segments. Expression extends to the other prosomal appendages, and by stage 10.2 (B-B'') it becomes restricted to the mesodermal tissue, with the exception of the chelicerae, where it is restricted to a small group of cells at the base (yellow arrow). This pattern remains unchanged at stage 12 (C-C''). Two additional expression domains were observed at stage 10.2 (B-B''), in the pre-cheliceral region (black arrowhead) and in the last pair of opisthosomal organs (red arrowhead). At stage 12 (C-C''), expression in the pre-cheliceral region is still present (black arrowhead), and faint expression was observed in all opisthosomal appendages (red arrowheads) and in the expanding dorsal tissue. *Pt-SoxE2* ISH staining (D-G'') revealed expression in the prosomal appendages similar to that of *Pt-SoxE1*, with the exception of the chelicerae, where expression extends to the tip (red arrow). At stage 8.2 (D), *Pt-SoxE2* is expressed in two clusters of cells in the pre-cheliceral region (yellow arrowhead), which starts fading at stage 9.1 (E) and is no longer visible during later stages (F and G). Specific and strong expression can be seen in all opisthosomal appendages (F', G' and G''; red arrowheads). Faint expression is also present in small clusters of ventral cells in the opisthosoma (black arrows). Anterior is to the left in all images. Ch, chelicerae; Pp, pedipalps; L1-L4, walking legs 1-4; O1-5, opisthosomal segment 1-5.

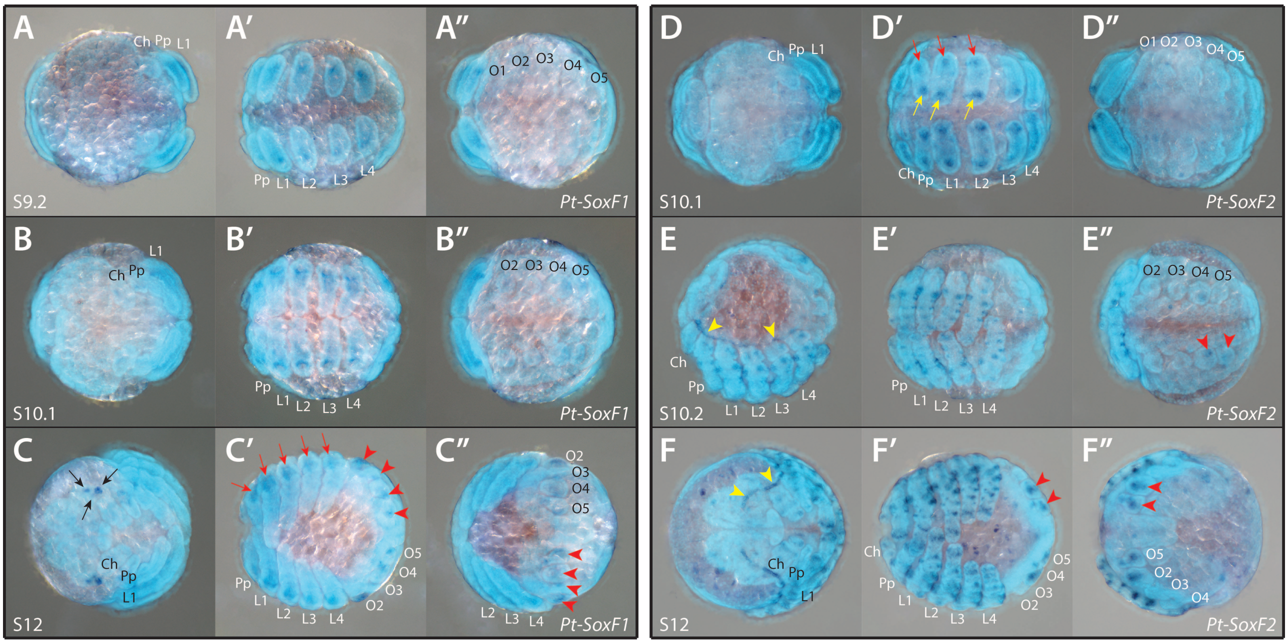


**Figure S9. Expression patterns of *P. tepidariorum SoxF* genes**

Faint expression of *Pt-SoxF1* is first visible at stage 9.2 (A-A'') in the prosomal appendages. This pattern remains unchanged up to stage 12 (B-B"), when it becomes restricted to the base of the legs (red arrows). At stage 12 (C-C''), additional expression domains arise in the lateral eye primordia (black arrows) and in the opisthosomal appendages (red arrowheads). *Pt-SoxF2* is also expressed at the tips (yellow arrows) and base (red arrows) of the prosomal appendages, starting at stage 10.1 (D-D"). This pattern is more complex at later stages and becomes restricted to several small groups of cells that possibly belong to the developing PNS (E' and F'). At stage 10.2 (E-E''), *Pt-SoxF2* expression appears along the edge of the prosomal region of the germ band, from the cheliceral to the L4 segments (yellow arrowheads). This domain extends to the pre-cheliceral region at stage 12 (F-F''), in a pattern that seems to follow the edge of the growing non-neurogenic ectoderm (yellow arrowheads). Lastly, *Pt-SoxF2* is also expressed in the last two pairs of opisthosomal appendages – the presumptive spinnerets (red arrowheads). Anterior is to the left in all images. Ch, chelicerae; Pp, pedipalps; L1-L4, walking legs 1-4; O1-5, opisthosomal segment 1-5.

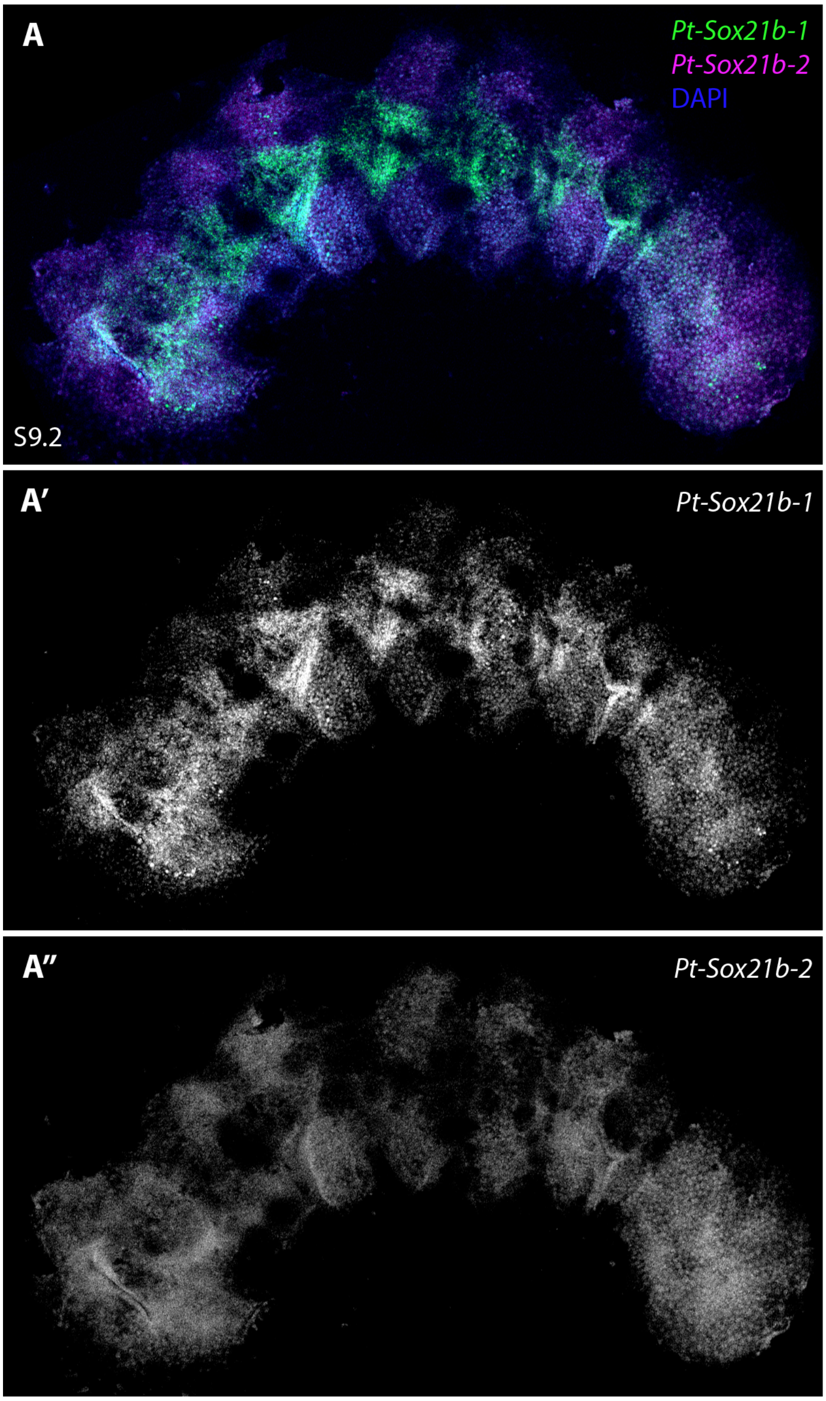


**Figure S10. Expression of *Pt-Sox21b-1* and *Sox21b-2***

At stage 9.1 *Pt-Sox21b-1* is expressed in the cephalic lobe, segmentally in the developing neuroectoderm and the SAZ (A, A'). *Pt-Sox21b-2* expression is limited to a faint signal in the developing limbs, parts of the neuroectoderm, as well as the cephalic lobe and the SAZ (A''). *Pt-Sox21-b1* and *Pt-Sox21b-2* expression appear to largely overlap in the cephalic lobe and the SAZ, whereas the expression overlap in the developing neuroectoderm is limited to the base of the developing limbs (A). Anterior is to the left in all images. Embryo is flat mounted. SAZ, segment addition zone; Ch, chelicerae; Pp, pedipalps; L1-L4, walking legs 1-4; O1, opisthosomal segment 1.

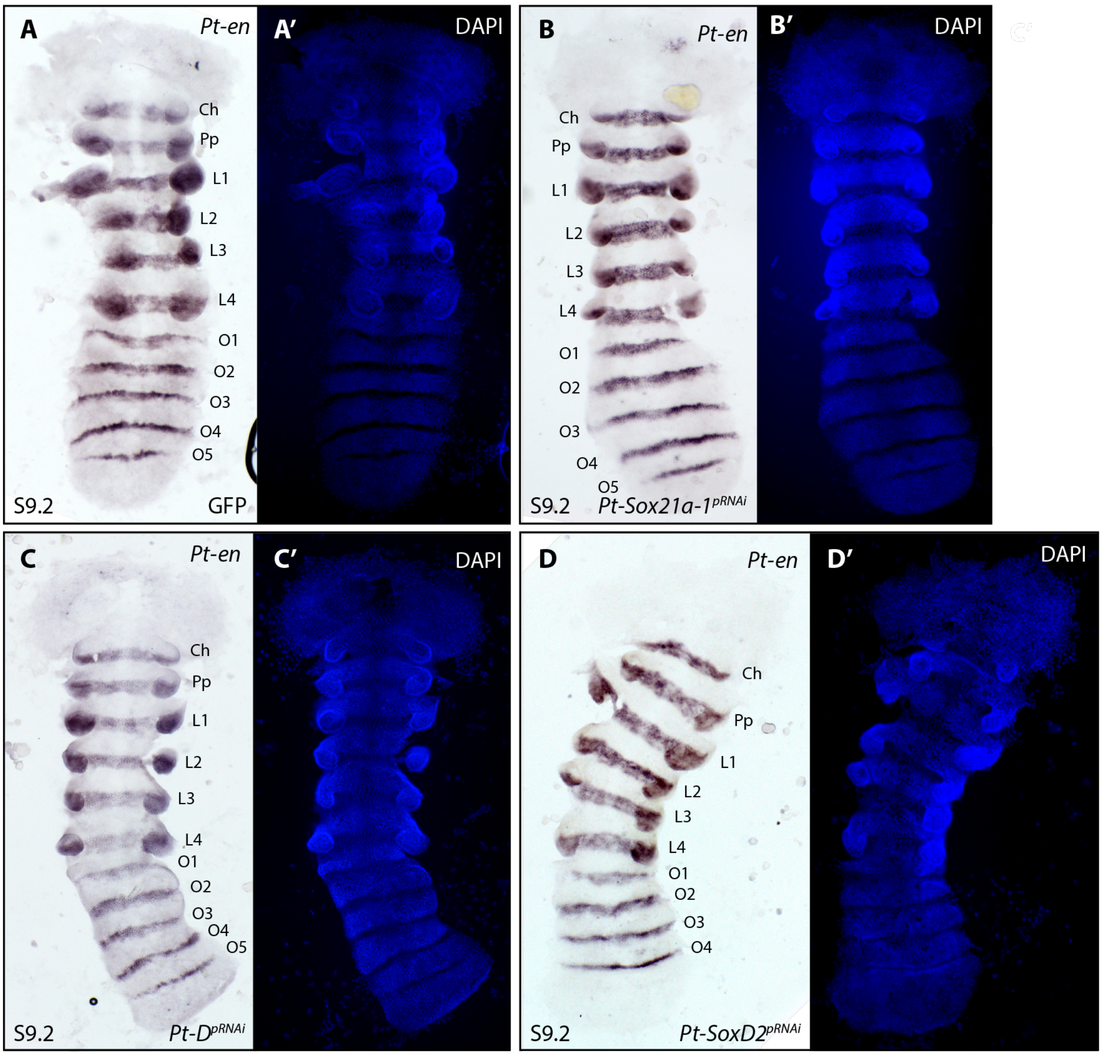


**Figure S11. Summary of *Pt-Sox21a-1*, *Pt-SoxD-2*, *Pt-D* parental RNAi**

At stage 9.2 *Pt-en* is expressed in all prosomal (Ch-L4) as well as opisthosomal segments (O1-O4/5) (A-D). All embryos displayed (A-D) were taken from 3rd cocoons: (A) shows a GFP control embryo, (B) shows a *Pt-Sox21a-1* knockdown embryo, (C) shows a *Pt-D* knockdown embryo and (D) shows a *Pt-SoxD2* knockdown embryo. All embryos (A-D) show normal development, no phenotypes were detected (detailed description in the text). Images (A', B', C',D') show the respective DAPI images. Anterior is at the top. All embryos are flat mounted. Ch, cheliceral segment, Pp, pedipalpal segment; L1-L4, walking leg segment 1-4 segments; O1-O5, opisthosomal segment 1-5

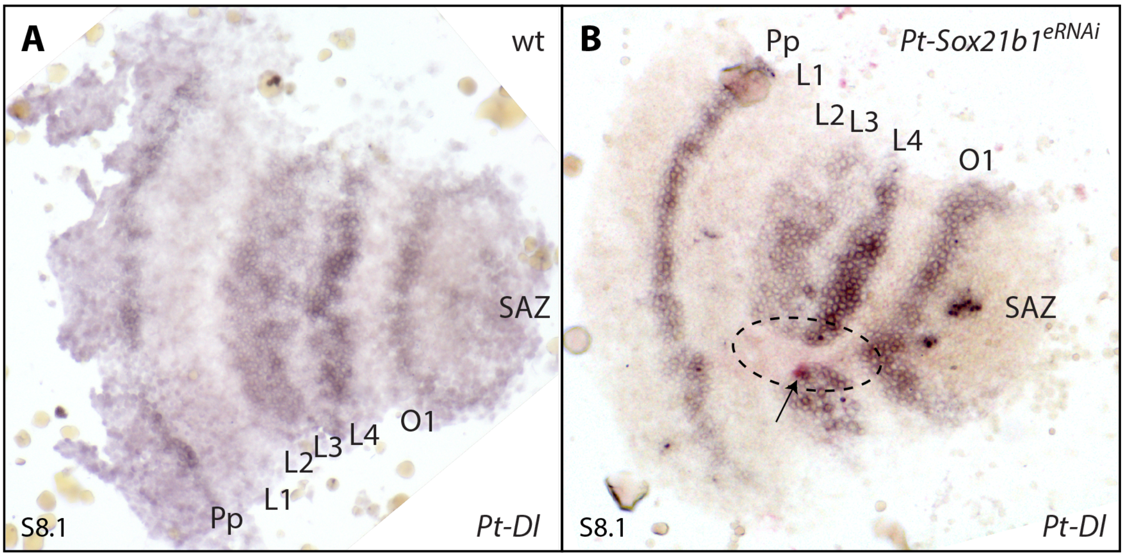


**Figure S12. Effect of *Pt-Sox21b-1* eRNAi knockdown on *Pt-Dl* expression**

At stage 8.1 (A), *Pt-Dl* is expressed in a stripe in the anterior SAZ, in the prosomal segments L2-L4 and the pedipalpal segment. At stage 8.1, *Sox21b-1* knockdown in the prosomal segments leads to a loss of *Pt-Dl* expression, as well as a coalescence of the surrounding tissue (B). Anterior is to the left. All embryos are flat mounted. Black arrow points at knockdown clone marked by pink staining; area of *Pt-Sox21b-1* knockdown is highlighted by dashed lines. Pp, presumptive pedipalpal segment; L1-L4, presumptive L1-L4 segments; O1, presumptive O1 segment; O2, presumptive O2 segment; SAZ, segment addition zone.
